# The Dynamic Conformational Landscapes of the Protein Methyltransferase SETD8

**DOI:** 10.1101/438994

**Authors:** Shi Chen, Rafal P. Wiewiora, Fanwang Meng, Nicolas Babault, Anqi Ma, Wenyu Yu, Kun Qian, Hao Hu, Hua Zou, Junyi Wang, Shijie Fan, Gil Blum, Fabio Pittella-Silva, Kyle A. Beauchamp, Wolfram Tempel, Hualiang Jiang, Kaixian Chen, Robert Skene, Y. George Zheng, Peter J. Brown, Jian Jin, Cheng Luo, John D. Chodera, Minkui Luo

## Abstract

Elucidating conformational heterogeneity of proteins is essential for understanding protein functions and developing exogenous ligands for chemical perturbation. While structural biology methods can provide atomic details of static protein structures, these approaches cannot in general resolve less populated, functionally relevant conformations and uncover conformational kinetics. Here we demonstrate a new paradigm for illuminating dynamic conformational landscapes of target proteins. SETD8 (Pr-SET7/SET8/KMT5A) is a biologically relevant protein lysine methyltransferase for *in vivo* monomethylation of histone H4 lysine 20 and nonhistone targets. Utilizing covalent chemical inhibitors and depleting native ligands to trap hidden high-energy conformational states, we obtained diverse novel X-ray structures of SETD8. These structures were used to seed massively distributed molecular simulations that generated six milliseconds of trajectory data of SETD8 in the presence or absence of its cofactor. We used an automated machine learning approach to reveal slow conformational motions and thus distinct conformational states of SETD8, and validated the resulting dynamic conformational landscapes with multiple biophysical methods. The resulting models provide unprecedented mechanistic insight into how protein dynamics plays a role in SAM binding and thus catalysis, and how this function can be modulated by diverse cancer-associated mutants. These findings set up the foundation for revealing enzymatic mechanisms and developing inhibitors in the context of conformational landscapes of target proteins.

## Introduction

Proteins are not static, but exist as an ensemble of conformations in dynamic equilibrium^1^. Characterization of conformational heterogeneity can be an essential step towards interpreting function, understanding pathogenicity, and exploiting pharmacological perturbation of target proteins^2–4^. Conventional efforts to map functionally relevant conformations rely on biophysical techniques such as X-ray crystallography5, nuclear magnetic resonance (NMR)^6^, and cryo-electron microscopy^7^, which provide static snapshots of highly-populated conformational states. While complementary techniques such as relaxation-dispersion NMR can resolve a limited number of low-population states, they are incapable of providing detailed structural information8. By contrast, molecular simulations provide atomistic detail---a prerequisite to structure-guided rational ligand design---and insight into relevant conformational transitions^1^. The emergence of Markov state models (MSMs) has shown the power of massively distributed molecular simulations in resolving complex kinetic landscapes of proteins^9,10^. By integrating simulation datasets with MSMs, functionally relevant conformational dynamics as well as atomistic details can be extracted^10^. Recently, MSMs have been used to identify key intermediates for enzyme activation^11,12^ and allosteric modulation^13^. However, these approaches are limited by the number of seed structures and timescales accessible by molecular simulations (generally microseconds for one structure) relative to the reality of complicated conformational transitions (up to milliseconds for multiple structures)^14^. To overcome the limitations of individual techniques, we envisioned an integrated approach that combines simulation with experiment to characterize conformational landscapes of enzymes and elucidate their functions with the consideration of dynamic conformations.

Protein lysine methyltransferases (PKMTs) comprise a subfamily of posttranslational modifying enzymes that transfer a methyl group from the cofactor *S*-adenosyl-L-methionine (SAM)^15^. PKMTs play epigenetic roles in gene transcription, cellular pluripotency, and organ development^16,17^. Their dysregulation has been implicated in neurological disorders and cancers^18,19^. SETD8 (SET8/Pr-SET7/KMT5A) is the sole PKMT annotated for monomethylation of histone H4 lysine 20 (H4K20me)^20,21^ and many non-histone targets such as the tumor suppressor p53 and the p53-stabilizing factor Numb^22,23^. Disruption of endogenous SETD8 leads to cell cycle arrest and chromatin decondensation, consistent with essential roles for SETD8 in transcriptional regulation and DNA damage response^24–26^. SETD8 has also been implicated in cancer invasiveness and metastasis^27^. High expression of SETD8 is associated with pediatric leukemia and its overall low survival rate^28^. As a result, there is enormous interest in elucidating functional roles of SETD8 in disease and developing pharmacological agents to perturb this target^29–31^.

Given the essential roles of conformational dynamics in enzymatic catalysis^1,32^ and our current limited knowledge of conformational landscapes of PKMTs, we envisioned leveraging an integrated experimental-computational approach to characterize dynamic conformational landscapes of SETD8 and its cancer-associated mutants with atomic resolution. To access previously-unseen, less-populated conformational states of SETD8 to seed massively parallel distributed molecular dynamics (MD) simulations, we envisioned trapping these conformations with small-molecule ligands. Here we solved four distinct crystal structures of SETD8 in alternative ligand-binding states with covalent SETD8 inhibitors and native ligands. With the aid of these new structures, we generated an aggregate of six milliseconds of explicit solvent MD simulation data for apo- and SAM-bound SETD8. Using a machine learning approach to select features and hyperparameters for MSMs via extensive cross-validation, we identified 24 kinetically distinct metastable conformational states of apo-SETD8 and determined how the conformational landscape is remodeled upon SAM binding. We then validated these conformational landscapes with stopped-flow kinetics and isothermal titration calorimetry by examining SAM binding, characterizing rationally-designed SETD8 variants with increased catalytic efficiency, and resolving multiple timescales associated with transitions among these conformers. The resulting model furnishes unprecedented key insights on how these dynamic conformations play a role in catalysis and how cancer-associated SETD8 mutants alter this process.

## Results

### Crystal structures of SETD8 associated with hidden conformations

To identify hidden high-energy conformational states of SETD8, we envisioned a strategy of trapping the associated conformers with small-molecule ligands. The development of high-affinity SETD8 inhibitors with canonical target-engagement modes is challenging^29^, and led us to exploit covalent inhibitors^31,33^. These compounds can overcome the high energy penalties associated with hidden high-energy conformers through the irreversible formation of energetically-favored inhibitor-SETD8 adducts. Our prior efforts led to the development of covalent inhibitors containing 2,4-diaminoquinazoline arylamide and multi-substituted quinone scaffolds by targeting Cys311^31,33^. Upon further optimization of these scaffolds, we identified MS4138 (**Inh1**) and SGSS05NS (**Inh2**)^34^, two structurally distinct covalent inhibitors with the desired potency against SETD8 (**Figures 1a**, **S9**). X-ray crystal structures of SETD8 were then solved in complex with **Inh1** and **Inh2**, respectively (**Figures 1b,c**, **S10**, **S11**). Notably, despite the overall structural similarity of the pre-SET, SET, and SET-I motifs, the **Inh1**- and **Inh2**-SETD8 binary complexes (**BC-Inh1**and **BC-Inh2**) differ from the SETD8-SAH-H4 ternary complex (**TC**)^35–37^ by the distinct conformations of their post-SET motifs. The post-SET motif of **TC**was characterized by its U-shaped topology with a double-kinked loop-helix-helix architecture, which appears to be optimally oriented for binding both SAM and a peptide substrate (**Figure 1c,d**)^35–37^. In comparison, **BC-Inh1**and **BC-Inh2** rotate their post-SET motifs by 140 ° and 60 °, respectively (**Figure 1d**). Moreover, the post-SET motifs of **BC-Inh1** and **BC-Inh2** adapt more extended configurations with a less structured loop and a singly-kinked helix, respectively (**Figure 1c,d**). Whereas multiple factors may influence the overall conformations, the formation of Cys311 adducts likely made the key contribution to the discovery of these hidden post-SET motif conformers.

**Figure 1.**
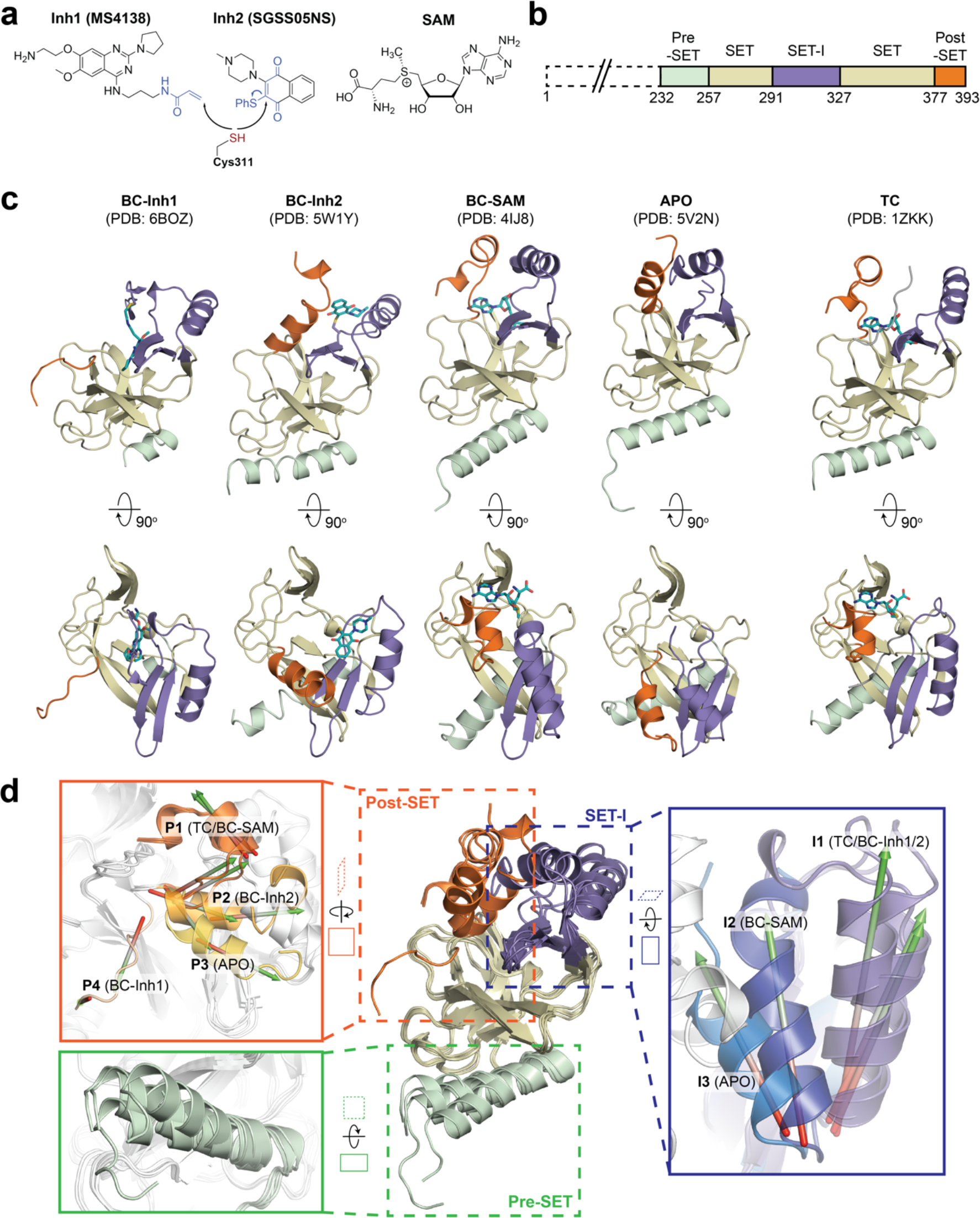
Diverse SETD8 conformations captured in altered ligand-binding states. **a**, Structures of SETD8 ligands involved in this work. Two covalent inhibitors targeting Cys311 (MS4138 as **Inh1**and SGSS05NS as **Inh2**) and the cofactor SAM were used as ligands to trap neo-conformations of SETD8. **b**, Domain topology of SETD8. Four functional motifs at SETD8’s catalytic domain are colored: pre-SET (light green), SET (dark yellow), SET-I (purple), and post-SET (orange). **c**, Cartoon representations of four neo-structures of SETD8 (**BC-Inh1**, **BC-Inh2**, **BC-SAM**, and **APO**) and a structure of a SETD8-SAH-H4 ternary complex (**TC**). These structures are shown in two orthogonal views with ligands, pre-SET, SET, SET-I, and post-SET colored in cyan, light green, dark yellow, purple, and orange, respectively. **d,**Superposition of five crystal structures highlighted with detailed views of post-SET, SET-I, and pre-SET motifs. The five X-ray structures reveal four distinct conformational states of the post-SET motif (**P1-4**) and three distinct conformational states of the SET-I motif (**I1-3**).

To reveal additional hidden conformers that are structurally distinct from **TC**, we also solved crystal structures of SETD8 upon depleting native ligands and obtained structures of the SAM-SETD8 binary complex (**BC-SAM**) and apo-SETD8 (**APO**) (**Figures 1c**, **S12**, **S13**). Strikingly, **BC-SAM** and **APO** differ from **TC** by their distinct SET-I motifs in the context of the otherwise similar SET-domain (**Figure 1d**). Furthermore, the post-SET motif of **APO** structurally resembles an intermediate state between **BC-Inh1** and **BC-Inh2** but is distinct from those of **BC-SAM** and **TC** (**Figure 1d**). In contrast to the structurally diverse SET-I (**I1-3**) and post-SET motifs (**P1-4**) in these structures, their pre-SET motifs show only slightly altered configuration (**Figure 1d**). The differences between these structures highlight the conformational plasticity of the SET-I and post-SET motifs. Collectively, these observations provide strong structural rationale for the existence of a highly dynamic conformational landscape of SETD8.

### Hidden conformations of apo-SETD8 revealed by structural chimeras

The **BC-SAM**, **BC-Inh1**, **BC-Inh2**, **APO**, and **TC**structures can be readily classified into three distinct SET-I configurations (**I1-3**) and four distinct post-SET configurations (**P1-4**) (**Figure 1d**). Given the relative independence between the SET-I and post-SET motifs, we expected that additional combinations of discrete motifs can represent yet-unobserved functionally relevant conformations of SETD8. We thus constructed putative “structural chimeras” of apo-SETD8 containing orthogonal **I1-3**and **P1-4**in a combinatorial (3×4) manner (**Figures 2a**, **S14**). Among the twelve structural chimeras as potential seeds for MD simulations, five were crystallographically-determined conformers (**BC-Inh1**, **BC-Inh2**, **BC-SAM**, **TC**with ligands removed, and **APO**), four were new structurally-chimeric conformers, and three were excluded because of obvious steric clashes (**Figures 2a**, **S15**).

**Figure 2.**
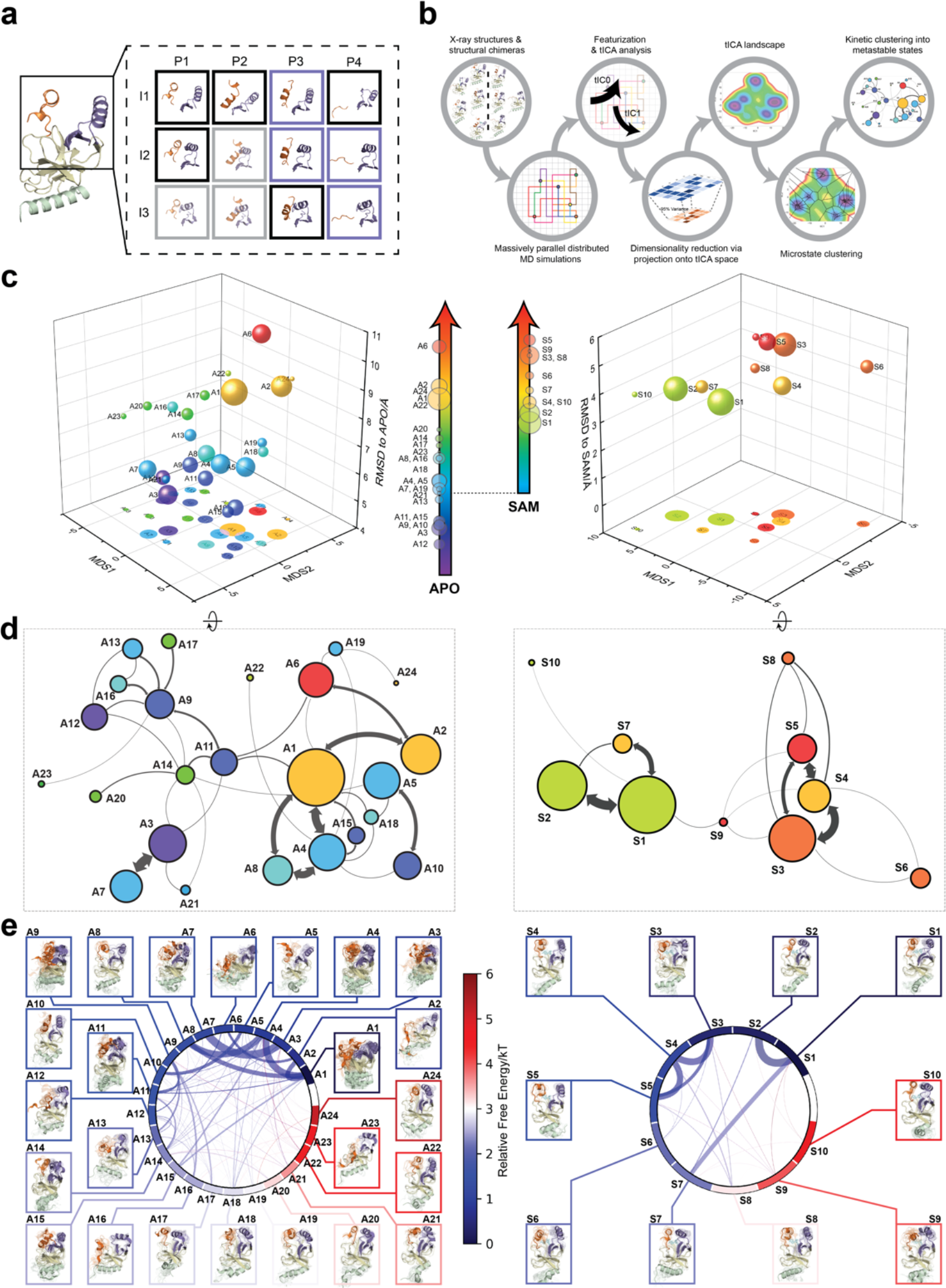
Markov state models and conformational landscapes of apo- and SAM-bound SETD8 constructed through diversely seeded, massively parallel molecular dynamics simulations. **a**, Combinatorial construction of structural domain chimeras using crystallographically-derived post-SET and SET-I conformations. Each conformer is boxed and color-coded with black for five X-ray-derived structures, blue for four putative structural chimeras included as seed structures for MD simulations, and grey for three structural chimeras excluded from MD simulations because of obvious steric clashes. **b**, Schematic workflow to construct dynamic conformational landscapes via MSM. The five X-ray structures and the four structural chimeras were used to seed massively parallel MD simulations on Folding@home (see Method). Markov state models were constructed from these MD simulation results to reveal the conformational landscape. **c–e**, Kinetically metastable conformations (macrostates) obtained from kinetically coupled microstates via Hidden Markov Model (HMM) analysis. The revealed dynamic conformational landscapes consist of 24 macrostates for apo-SETD8 (left panel) and 10 macrostates for SAM-bound SETD8 (right panel). **c**, Kinetic and structural separation of macrostates in a 3D scatterplot. The X, Y axes represent kinetic separation of macrostates with a log-inverse flux kinetic embedding method (see Methods). The Z axis reports RMSDs of each macrostate to **APO** (left) or **BC-SAM** (right). The relative population of each macrostate of apo- or SAM-bound SETD8 ensembles is proportional to the volume of each representative sphere. **d**, Cartoon depiction of macrostates in a 2D scatterplot. The relative positions of metastable conformations were derived via the log-inverse flux kinetic embedding (see Methods). The diameter of the corresponding circle in the 2D scatterplot is proportional to the diameter of the respective sphere in the 3D scatterplot above. Equilibrium kinetic fluxes larger than 7.14×10^2^ s^−1^ for apo- and 1.39×10^3^ s^−1^ for SAM-bound SETD8 are shown for interconversion kinetics with thickness of the connections proportional to fluxes between two macrostates. **e**, Chord diagrams and representative conformers of macrostates. The colors are encoded for the free energy of each macrostate relative to the lowest free energy of the macrostates. The equilibrium flux between two macrostates is proportional to thickness of respective arcs.

### Dynamic conformational landscape of apo-SETD8 via Markov state modeling from 5-ms MD simulation dataset

With seed conformations prepared as above, we envisioned illuminating the conformational landscape with massively distributed long-time MD simulations and resolving its kinetic features with Markov state models (MSMs) (**Figures 2b**, **S14**). We conducted approximately 500×1 µs explicit-solvent MD simulations from each seed and accumulated 5 milliseconds of aggregate data in 10 million conformational snapshots for apo- SETD8 (**Figures S16**, **Table S3**). To identify functionally relevant conformational states and their transitions, we built MSMs using a pipeline that employs machine learning and extensive hyperparameter optimization to identify slow degrees of freedom and structural and kinetic criteria to cluster conformational snapshots into discrete conformational states (**Figures S17-24**, **Tables S4**, **S5**)^38^. This approach identified 24 kinetically metastable conformations (macrostates) from an optimized, cross-validated set of 100 microstates (**Figures 2c**, **S25-30**, **Tables S6**, **S7**). These macrostates are remarkably diverse, spanning up to 10.5 Å Cα RMSD from **APO**. To visualize the kinetic relationships between functionally important conformations, dimensionality reduction was used to project the landscape into 2D while preserving log inverse fluxes between states (**Figure 2d**). The relative populations of these macrostates and their interconversion kinetics were calculated on the basis of their transition fluxes, resolving rare conformational states up to 6 kT in free energy (**Figure 2d,e**).

### The dynamic conformational landscape of SAM-bound SETD8

Given the success in constructing the dynamic conformational landscape of apo-SETD8, we applied the same strategy to SAM-bound SETD8. With the two crystal structures of SETD8 in complex with SAM (**BC-SAM**and **TC**) as the seed conformations, we conducted ~ 500×1 µs explicit-solvent MD simulations from each structure and accumulated 1 millisecond of aggregate data (2M snapshots) (**Figure S25**). The resulting MSM for SAM-bound SETD8 contained 10 kinetically metastable macrostates arising from 67 microstates (**Figure S31**, **Tables S8**, **S9**). Similar to those of apo- SETD8, the relative macrostate populations of SAM-bound SETD8 and their flux kinetics were computed and embedded into 3D/2D scatter plots and chord diagram (**Figure 2c,d,e**). The smaller number of metastable states identified for SAM-bound SETD8 is expected given that SAM binding restricts conformational accessibility.

**Figure 3.**
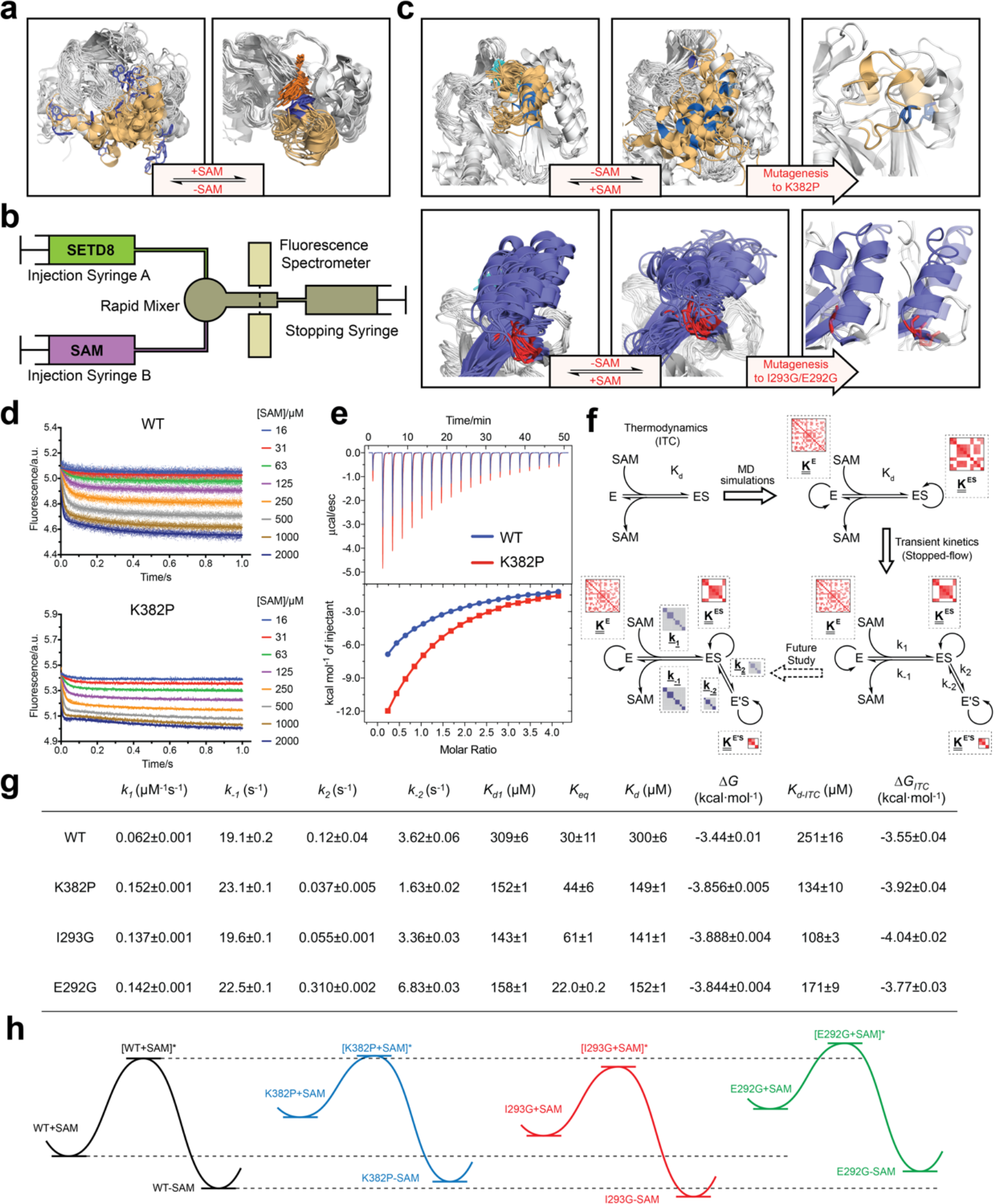
Biochemical characterization of gain-of-function mutations revealed by conformational landscapes of SETD8. **a**, Comparison of binding environments of Trp390 between apo and SAM-bound SETD8 in the context of their dynamic conformational landscapes. **b**, Illustration of rapid-quenching stopped-flow experiments. These experiments were conducted to trace fluorescence changes of Trp390 upon SAM binding. **c**, Comparison of the conformations of post-SET kink and SET-I helix between apo and SAM-bound SETD8 in the context of their dynamic conformational landscapes. Analysis of key structural motifs indicated K282P, I293G and E292G as potential gain-of-function variants. **d**, Fluorescence changes of wild-type and K382P SETD8 traced with a rapid-quenching stopped-flow instrument within 1 s upon SAM binding. **e**, SAM-binding ITC enthalpogram of wild-type and K382P SETD8. **f**, Stepwise SAM-binding of SETD8 in the integrative context of biochemical, biophysical, structural, and simulation data. ITC determines the thermodynamic constant of SAM binding by SETD8. MD simulations and MSM uncover metastable conformations and interconversion rates of apo- and SAM-bound SETD8 (*<u>K</u>*^apo^ and *<u>K</u>*^SAM^). Stopped-flow experiments revealed that SETD8 binds SAM via biphasic kinetics. Rate constants uncovered by stopped-flow experiments (*k*_1_, *k*_−1_, *k*_2_, *k*_-2_) represent macroscopic rates of SAM binding by SETD8 with multiple metastable conformations. The microscopic behavior of individual metastable states and corresponding rates 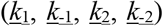 have not been resolved. Transition probability matrices (red) and microscopic rate constant matrices (blue) are shown as colored grids. A rigorous mathematical model for these processes is shown in **Figure S36**. **g**, Kinetic and thermodynamic constants of wild-type SETD8 and its mutants. For *k_1_*, *k_−1_*, *k_2_*, *k_−2_*, data are best fitting values ± standard error (s.e.) from KinTek. For *K_d-ITC_*, data are mean ± s.e. of at least 3 replicates. *K*_d1_, *K*_eq_, and *K*_d_ are calculated based on equations in online method. Uncertainties of *K*_d1_, *K*_eq_, *K*_d_, and ΔG are s.e. calculated by the propagation of uncertainties from individual rate constants and dissociation constants, respectively. **h**, Relative energy landscapes of apo- and SAM-bound SETD8 and its gain-of-function mutants.

### Experimental validation of the conformational landscapes of SETD8

Upon uncovering the dynamic conformational landscapes of apo- and SAM-bound SETD8, we were able to extract new structural information and designed experiments to further validate this model (**Figure 3**). Comparison of the conformational ensembles between apo- and SAM-bound SETD8 revealed that SAM binding dramatically alters the environment of Trp390 (**Figure 3a**, blue sticks), the sole tryptophan residue in the catalytic domain of SETD8. This residue is flexible and mainly solvent-exposed in apo-SETD8 conformational ensembles but restricted in a hydrophobic environment through SAM-mediated pi-pi stacking in SAM-bound SETD8 conformational ensembles (**Figure 3a**). Such environmental changes upon SAM binding are expected to quench fluorescence of Trp390^39^. To verify this prediction, we designed rapid-mixing stopped-flow kinetic experiments with 5 ms dead time and 0.1 ms resolution to track the fluorescence change of Trp390 upon SAM binding (**Figure 3b**). We observed SAM-dependent biphasic kinetics of the fluorescence decrease within 1 s with > 80% of the change occurring in the fast phase (0 – 0.1 s) (**Figure 3d**). In the context of the conformational landscape of apo-SETD8, we interpreted the major decrease in fluorescence intensity (fast-phase kinetics) as a consequence of the collective changes of Trp390 from the solvent-exposed hydrophilic environment in apo conformations to the hydrophobic environment in SAM-bound conformations (**Figure 3a,c**). In contrast, the minor decrease in fluorescence intensity (slow-phase kinetics) reflects the slow conformational changes of Trp390 in the SAM-bound SETD8 conformational ensembles (**Figure 3d**). With unsupervised global fitting to this two-step model, we obtained forward and reverse rate constants for the fast- and slow-phase kinetics, which are in agreement with conventional fitting to double exponential kinetics^40^ (**Figures 3d,f,g**, **S32**, **Table S10**). The *k*_−1_ value was also confirmed independently by rapid-mixing stopped-flow dilution of SAM-bound SETD8^41^ (“ES+E’S”, **Figure S33**, **Table S10**). Here the *k*_-1_/*k*_1_ ratio of 309±6 μM corresponds to the average SAM dissociation constant *K*_d1_ of apo-SETD8 conformers, which is consistent with independently determined ITC *K*_d_ of 251±16 μM (**Figures 3e,f**, **S34**). In contrast, the large *k*_-2_*/k*_2_ ratio of 30±11 suggests that the second phase corresponds to a slow equilibrium between ES and E’S with minimal contribution of E’S to the overall SAM dissociation constant *K*_d_ (**Figure 3e**). The conformational ensembles we identified for apo- and SAM-bound SETD8 demonstrate the statistical nature of its SAM-binding process. Therefore, the observed fluorescence changes and herein determined macroscopic kinetic constants represent an ensemble-weighted average of microscopic behaviors of all species that exist in the solution. A rigorous mathematical description of microscopic kinetics of SAM binding was thus obtained under the consideration of interconversion of the metastable conformational states of apo- and SAM-bound SETD8 (**Figure S36**).

We then proposed to confirm our understanding of functionally-relevant conformations and their thermodynamics by identifying SETD8 variants with increased affinity for SAM. We uncovered a collection of characteristic kink motifs around Lys382 in the post-SET motif of SAM-bound SETD8 conformational ensembles (**Figure 3c**), while this region is less structured in apo-SETD8 conformational ensembles. We hypothesized that a proline mutation (K382P) could better stabilize the conformational ensembles of SAM-bound SETD8 than apo-SETD8 (**Figure 3c,h**). We also identified a characteristic α-helix in the SET-I motif, which adapts flexible and diverse configurations in apo ensembles but constrained and structurally distorted configurations in SAM-bound ensembles (**Figure 3c**). We proposed that the replacement of I293 or E292 adjacent to the α-helix with a flexible glycine should relax this distortion to better stabilize SAM-bound ensembles (**Figure 3c,h**). We therefore characterized the SAM-binding kinetics and affinities of K382P, I293G, and E292G variants of SETD8 with stopped-flow kinetics and ITC (**Figures 3c,d,e,f**, **S32-34**). While exhibiting biphasic kinetics similar to that of wild-type SETD8, the stopped-flow mixing experiment revealed the three variants showed a significant two-fold decrease of *K*_d,SAM_ (**Figure 3d,e**). The stopped-flow data further revealed that the two-fold change of *K*_d,SAM_ mainly arises from increased SAM-binding rates *k*_1_ with relatively unchanged *k*_-1_ (**Figure 3g**). These results are consistent with independently-determined *K*_d_ and *k*_-1_ from ITC and stopped-flow dilution, respectively (**Figures 3e,f**, **S33**, **S34**, **Table S10**). Collectively, these observations confirm the robustness of our conformational landscape model for apo- and SAM-bound SETD8.

### Effects of key simulation parameters on construction of conformational landscapes

We systematically investigated how the choices of seed structures and simulation time---key computational parameters---influence microstate discovery and quality of conformational landscapes of SETD8 (**Figure 4**). The simulations of apo-SETD8 initiated from any single X-ray structure (**BC-Inh1**, **BC-Inh2**, **BC-SAM**, **APO**, or **TC**in **Figure 1c**) only reveal a partial conformational landscape (28–61% microstate coverage, **Figure 4a**). To achieve >90% microstate coverage, at least two crystal structures---**BC-SAM**in combination with either **BC-Inh1**or **BC-Inh2**---must be included (**Figure 4a**). If three crystal structures are included, **BC-SAM**in combination with **TC**and **APO**can provide >90% coverage (**Figure 4a**). In terms of the structural motifs (**I1-3**or **P1-4**, **Figures 1d**, **2a**), simulations originating from the SET-I motif **I1**, **I2**, or **I3**alone led to the discovery of 69, 58, or 39 of the 100 microstates, respectively (**Figure 4b**, **Table S15**). The combination of **I1**and **I2**is sufficient to cover all 100 microstates, arguing for the redundant character of **I3**. For the post-SET motif, any combination of two post-SET configurations except **P2**–**P3**leads to >90 microstate coverage (**Figure 4b**, **Table S15**). These findings are in agreement with the key requirement of structural motif conformations **I1**(equivalent to **BC-Inh1**, **BC-Inh2**, or **TC**), **I2**(equivalent to **BC-SAM**), and any two of **P1**–**4**except **P2**–**P3**(*e.g.* **P1**–**P3**is equivalent to the combination of **APO**with **BC-SAM**or **TC**) to achieve >90% microstate coverage. For SAM-bound SETD8, the seed conformations derived from **BC-SAM**and **TC**structures contribute 31 and 38 of 67 microstates (**Figure 4c,d**, **Table S14**). These findings argue for the importance of using multiple structures to construct the landscape within achievable computer time. The seed conformations prepared from ligand-trapped SETD8 structures are essential to discovering the complete conformational landscapes of SETD8.

For simulation time, we observed that the fewer seed conformations of apo-SETD8 were employed, the more computing power (the product between the number of simulation trajectories and the time length per trajectory) was required to reach a comparable level of microstate coverage (**Figure 4e**, **Tables S16**, **S17**). When computing power is fixed, comparable microstate coverages of apo- and SAM-bound SETD8 can be obtained by running either multiple short trajectories or few long trajectories (**Figures 4f**, **S38**). The current simulation time (5 ms for apo- SETD8 and 1 ms for SAM-bound SETD8) provides 2–10-fold redundant computing power to map the conformational landscapes of SETD8.

**Figure 4.**
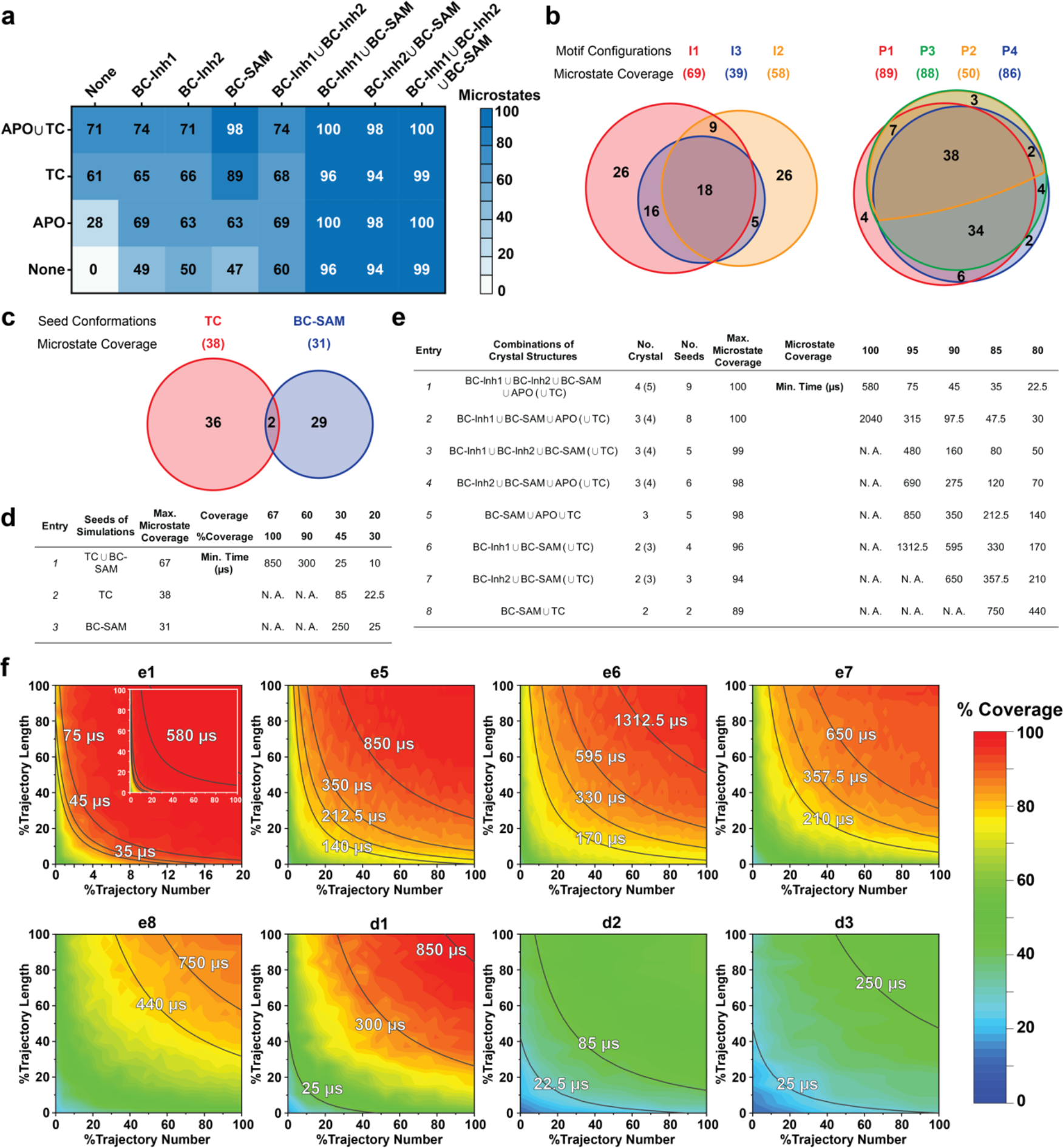
Evaluation of key simulation parameters of massively distributed molecular simulations. **a–b**, Robustness of simulations of apo-SETD8: **a**, Heat map for the coverage of the 100 microstates with all combinations of the crystal structures (**BC-Inh1, BC-Inh2, BC-SAM, APO, and TC**) as seed conformations; **b**, Venn diagrams of the coverage of the 100 microstates with all conformational combinations of SET-I and post-SET motifs (**I1-3**and **P1-4**) as seed structures for MD simulations. **c–d**, Robustness of simulations of SAM-bound SETD8: **c**, Venn diagram of the coverage of the 67 microstates with **TC**, **BC-SAM** or both as seed structures for MD simulation; **d**, Minimal time required by MD simulations to reach certain coverage of the 67 microstates with representative combinations of seed structures. **e**, Minimal time required by MD simulations to reach certain coverage of the 100 microstates of apo-SETD8 with representative combinations of seed structures. **f**, Contour map of microstate coverage at various combinations of trajectory lengths and numbers as percentage of the maximal trajectory length and number of MD simulations. The seed structures of each panel are listed as the simulation entries e1, e5–8 for apo-SETD8, and d1–3 for SAM-bound SETD8. Each curve corresponds to the aggregation of specific simulation time.

### Functionally relevant conformations in the dynamic landscapes of apo- and SAM-bound SETD8

After validating the conformational landscapes of apo- and SAM-bound SETD8, we explored the dynamic details of these landscapes with the focus on the connectivity and equilibrium fluxes between kinetically metastable macrostates (henceforth referred to as the “network”). When projected into two dimensions, the conformational landscape of apo-SETD8 takes the form of a dumbbell-like shape containing two lobes, each composed of about 12 macrostates primarily connected via a single hub-like central macrostate A11 (**Figures 2d**, **5**, **Table S7**). The conformational landscape also consists of other multiply-connected macrostates, including A1–A4, A9, and A14, as characterized by their rapid kinetic interconversion with multiple other macrostates (**Figure 2d,e**). Most low-populated macrostates (A17–A24) appear as satellite macrostates in the periphery of the network with few high-flux channels of interconversion to other macrostates (**Figure 2d,e**). The remaining states were classified as basin-like macrostates including (A5, A10), A7, A8, (A12, A13, A16) and A15, because these macrostates are highly populated and either relatively isolated or appear in tightly interconnected but globally isolated groups.

**Figure 5.**
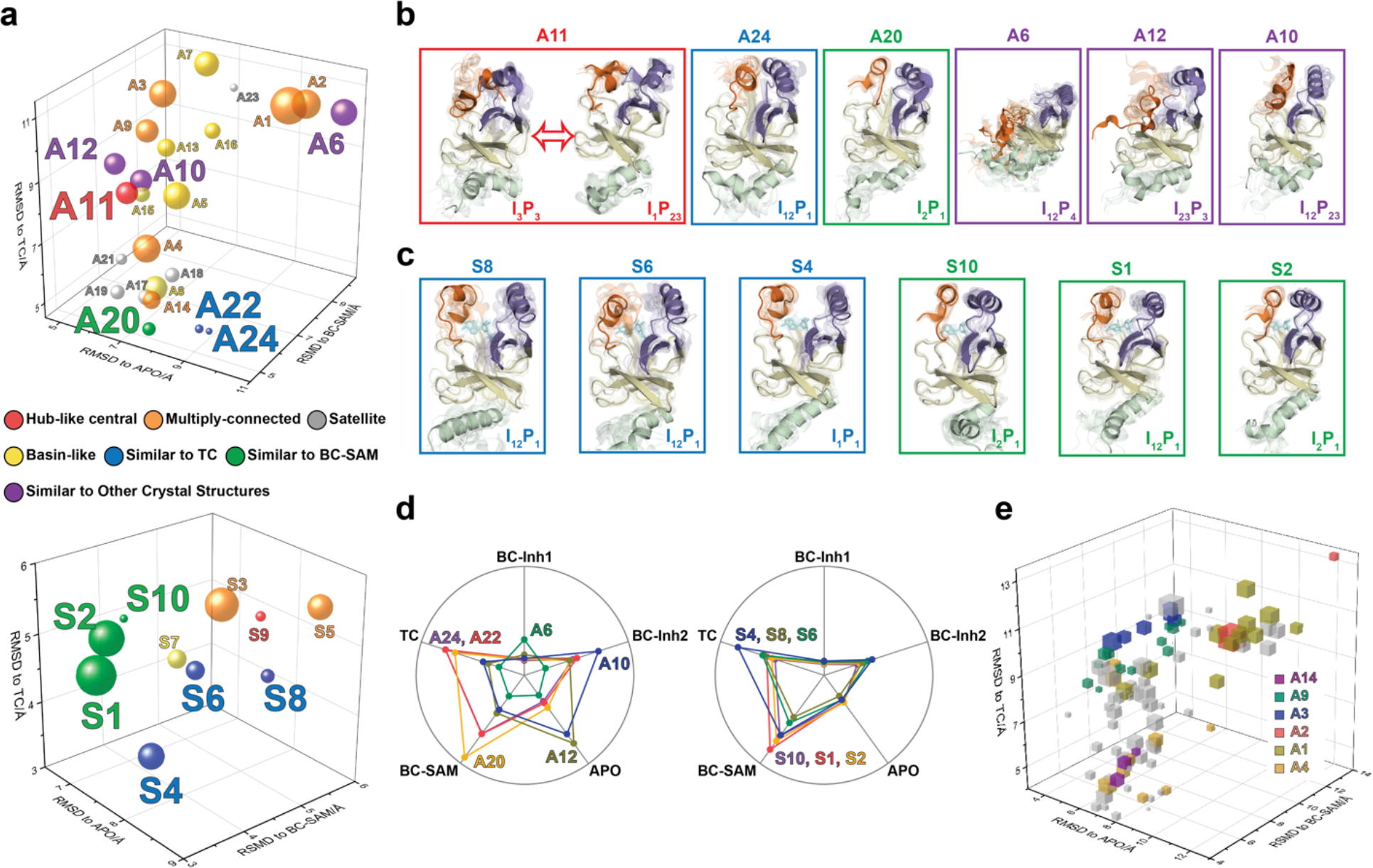
Functional annotation of the dynamic conformational landscapes of SETD8. **a**, 3D scatterplots of the 24 macrostates of apo-SETD8 landscape and 10 macrostates of SAM-bound SETD8 landscape in the coordinates of RMSDs relative to **APO**, **BC-SAM**, and **TC**. Volume of each sphere is proportional to the relative population of the corresponding macrostate in the context of the 24 macrostates for apo-SETD8 or the 10 macrostates for SAM-bound SETD8. The RMSD of each macrostate is the average of its microstates weighted with their intra-macrostate population. The RMSD of each microstate is the average of the top 10 frames most closely related to the clustering center of the microstate. The feature of each macrostate is annotated in color. **b**, **c** Cartoons of representative conformations of key macrostates in the apo-SETD8 landscape and the SAM-bound SETD8 landscape, respectively. Structural annotations are shown in bottom right of each conformation. **d**, Radar chart of representative macrostates of apo (left) and SAM-bound (right) landscapes in reference to the five crystal structures. Distances between dots and cycle centers are proportional to the reciprocal values of RMSDs of macrostates relative to the crystal structures. **e**, 3D scattering plot of 100 microstates of the apo landscape in the coordinates of RMSDs to **APO**, **BC-SAM**, and **TC**. Volume of each cube is proportional to the relative population of the corresponding microstate in the context of the 100 microstates. Microstates clustered in intermediate-like macrostates are highlighted in colors. Structural diversity of microstates within individual macrostates indicates that each intermediate-like state contains multiple structurally distinct but readily interconvertible microstates.

The hub-like macrostate A11 consists of two structurally-distinct microstates with comparable populations (**Figures 2d**, **5a**). One microstate structurally resembles the conformation of **APO** (I_3_P_3_), while the other microstate represents a conformer with the I_1_P_23_ feature for its SET-I and post-SET motifs (**Figure 5b**, **Table S6**). Rapid conformational interconversions within A11 is consistent with its hub-like character, centered between the two lobes of the dumbbell-like network. Interestingly, macrostates kinetically adjacent to A11 have structurally similar SET-I motifs within each lobe but distinct SET-I motifs between the two lobes (**I2**~**3** for the left and **I1**~**2** for the right) (**Figures 2d**, **5b**). Therefore, A11 is a transition-type state essential for the conformational fluxes of the macrostates between the two lobes, involved in a key step of conformational changes of the SET-I motif between **I1**~**2** and **I2**~**3**.

The intermediate-like macrostates A1–A4, A9, and A14 each contains multiple structurally distinct but kinetically associated microstates (**Figures 2d**, **5a**,**b**). The satellite macrostates A17–A24 are less populated and more structurally homogeneous (**Figures 2d**, **5a,b**). Conformers in the macrostates A22, A24 and A20 are structurally similar to **TC** and **BC-SAM** with slightly different but well-defined SAM-binding pockets, suggesting minimal conformational reorganization of A22, A24, and A20 is required to accommodate the cofactor (**Figure 5a,b,e**). Interestingly, A22 and A24, whose overall structures are similar to each other (**TC**-like), rarely interconvert in the apo landscape (**Figure 2d**). In contrast, the basin-like macrostates (A5, A10), A7, A8, (A12, A13, A16) and A15 do not contain a well-defined SAM-binding pocket (**Figure 5a,b,e**). Here the conformers in macrostate A12 are similar to **APO**, the conformers in the macrostate A6 are similar to **BC-Inh1**, and the conformers in the macrostates A10 are similar to **BC-Inh2**(**Figure 5d**). The structural similarity between the simulated conformers and **BC-Inh1/2**strongly argue that the two covalent inhibitors successfully trapped key hidden conformers of apo-SETD8.

Similar to that of apo-SETD8, the interconversion network of the macrostates of SAM-bound SETD8 also displays a dumbbell-like shape with S9 as the hub-like state connecting the two lobes of the network (**Figures 2d**, **5a**). The macrostates S1 and S3–S5 are multi-connected states; S6, S8, and S10 are satellite-like states; S2 and S7 are basin-like states (**Figure 5a,b**). Notably, the complexity of the overall conformational landscape of SAM-bound SETD8 is dramatically reduced in comparison with those of apo-SETD8 (**Figures 2d**, **5a**). The conformers in S1, S2, and S10 are structurally similar to those of A20, as well as **BC-SAM**; the conformers in S4, S6, and S8 are structurally similar to those in A22 and A24, as well as **TC** (**Figure 5c,d**). The structural similarities between these apo and SAM-bound macrostates suggest possible pathways for connecting the two conformational landscapes upon SAM binding.

**Figure 6.**
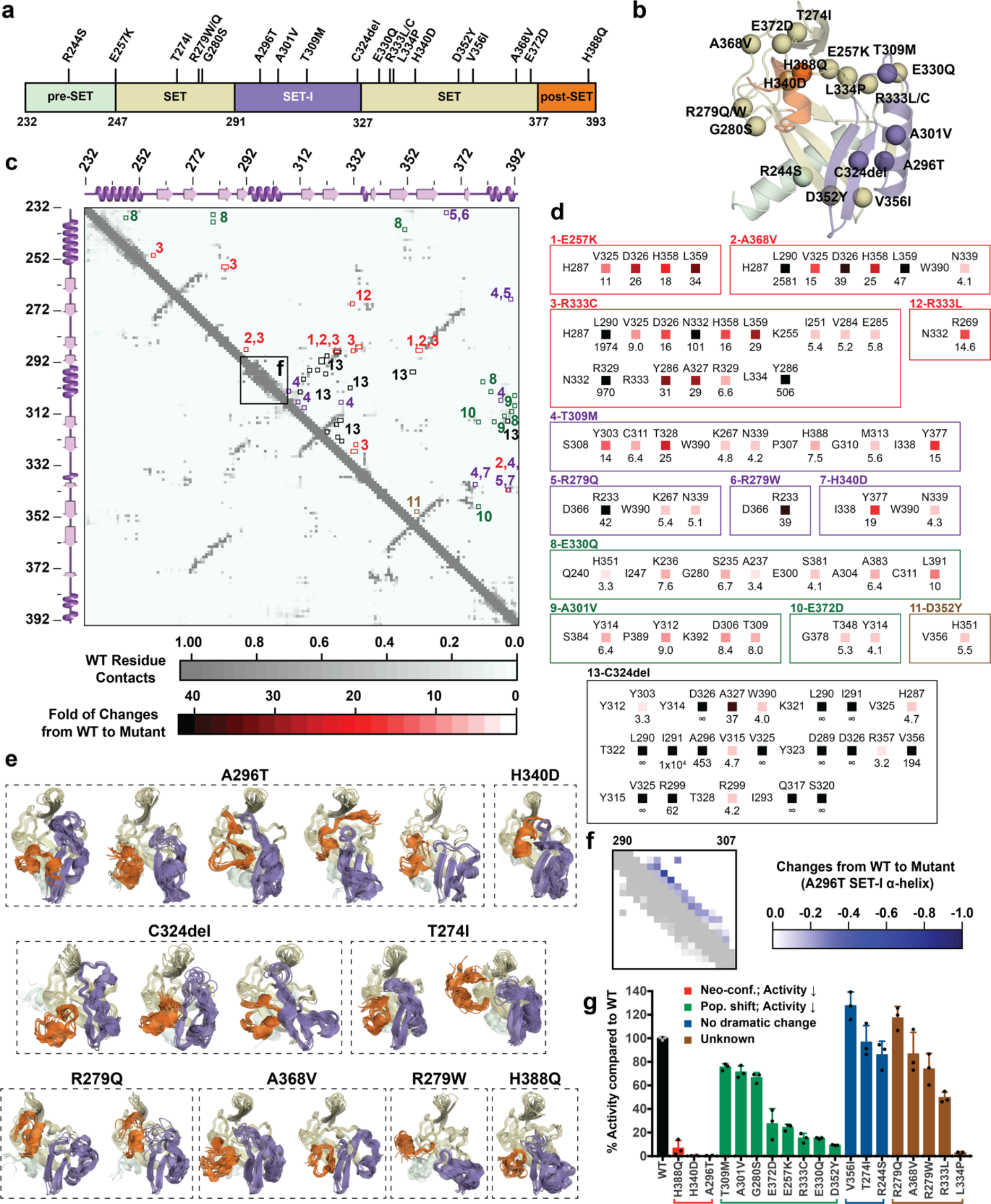
Computational and experimental characterization of cancer-associated SETD8 mutants. **a**, Cancer-associated mutations in the catalytic domain of SETD8 examined in this work. **b**, Cartoon representations of **TC** with cancer-associated SETD8 mutations highlighted. **c**, Differential residue-contact maps of cancer-associated SETD8 mutants in reference to wild type apo-SETD8 (grey). **d**, Representative contacts in the differential residue-contact maps of cancer-associated SETD8 mutants. The contacts of SETD8 mutants with >3-fold gain of contact fraction relative to wild-type SETD8 are listed and color-coded according to the increased magnitude of the contact fraction. **e**, Cartoon representations of neo-conformations revealed by simulations of SETD8 mutants. **f**, Differential residue-contact maps of the structurally relaxed α-helix at the SET-I motif of SETD8 A296T mutant. Decrease of contact fraction of SETD8 mutants relative to wild-type SETD8 is colored in blue. **g**, Enzymatic activities of wild-type and mutated SETD8 determined by an *in vitro* radiometric assay with H4K20 peptide substrate. Here SETD8 mutants are categorized as the following: red, uncovered neo-conformations (Neo-conf.) with > 90% loss of methyltransferase activity; green, populated inactive conformations (Pop. shift) with partially abolished methyltransferase activity; blue, no dramatic change of differential contact maps with comparable methyltransferase activity with wild-type SETD8; brown, unknown relationship between differential contact maps and methyltransferase activities. Data are mean ± standard deviation (s.d.) of 3 replicates.

### Characterization of cancer-associated SETD8 mutants

Sequences from tumor samples retrieved from cBioPortal^42–44^ contain two dozen point mutations in the catalytic domain of SETD8 (**Figure 6a,b**, **Table S11**). We expect that some of these mutations perturb SETD8 function. Because of conformational heterogeneity, it has historically been challenging for ***in silico*** approaches to annotate how mutations---in particular those structurally remote from functional sites---affect a target protein on the basis of its static structure(s) ^45–47^. Here, we envisioned addressing this challenge with the aid of the dynamic conformational landscapes of SETD8. To characterize mutations remote from catalytic sites (around 80% of known mutations), 40 independent microsecond-long MD simulations for each of the cancer-associated apo-SETD8 mutants were conducted with seed structures prepared from one ternary complex (**TC**) conformer. We then constructed a differential residue-contact map for each variant (**Figure 6c,d**) and extracted snapshots representing most dramatic conformational deviations from the wild type conformational ensembles (**Figure 6e**). Remarkably, even with modest simulation time, several cancer-associated mutants displayed neo-conformations that were not observed in the 5 ms wild-type dataset and cannot be predicted from static X-ray crystal structures. Strikingly, all of the neo-conformations display distinct reorganizations at the SET-I motif (**Figure 6e**). For instance, a single point mutation A296T, ~16 Å remote from the active site, yields five distinct neo-conformations (**Figure 6e**). In addition, relative to wild-type apo-SETD8, this mutant populates several conformations with a structurally relaxed α-helix at the SET-I motif (**Figure 6e**). C324del, ~20 Å from the SET-I motif, is associated with three neo-conformations and displays the most dramatic changes in the differential contact map (**Figure 6d**, panel 13). The remote H340D mutation is associated with one neo-conformation as well as more populated conformations containing spatially compressed active sites (**Figure 6d**, panel 7; **6e**). Using *in vitro* radiometric assays, the A296T and H340D mutants were characterized by loss of the methyltransferase activity on H4K20 peptide substrate (**Figure 6g**). The failure to purify recombinant C324del also supports the impact of this deletion on SETD8 function. H388Q, which mutates a histidine involved in substrate binding, is also associated with neo-conformations as well as loss of the methyltransferase activity (**Figure 6e,g**). These observations provide potential molecular rationale for how remote mutations can alter the active sites and the SET-I motif---and hence catalysis---via modulating the conformational landscape. Exceptions are T274I, R279W, R279Q, and A368V, which yielded neo-conformations but showed activity comparable to wild-type SETD8 (**Figure 6e,g**).

The differential residue-contact maps further revealed that remote mutations can alter conformational landscapes by altering populations of pre-existing conformations (**Figure 6c,d**). For instance, E257K, G280S, A301V, T309M, E330Q, D352Y mutations populate conformations containing spatially compressed active sites (**Figure S37**); E372D populates conformations containing a constrained post-SET motif; R333C populates conformations with reorganized SET motifs adjacent to the peptide binding pocket. All of these mutations showed partial loss of methyltransferase activity (**Figure 6g**). Notably, these structural alterations are often remote from the corresponding mutation sites (**Figure 6b**). In contrast, R244S, T274I, and V356I showed no significant conformational change on the basis of their differential contact maps, consistent with their comparable methyltransferase activity to wild type SETD8 (**Figure 6g**). Likely due to insufficient simulation time (40×1 μs/mutant), R333L and L334P variants, characterized by partial-to-complete loss of the methyltransferase activity (**Figure 6g**), showed similar conformational landscapes to that of wild-type apo-SETD8. Exploring conformational landscapes is thus an effective strategy to reveal structural alterations associated with majority of remote-site mutations for functional annotation.

## Discussion

Here we have demonstrated that tight integration of structural determination---using covalent probes and multiple ligand-binding states to trap hidden conformations (**Figure 1**)---with massively distributed molecular simulations and the powerful framework of Markov state models (**Figure 2b**) can provide unprecedented insights into the detailed conformational dynamics of an enzyme. The current work demonstrates the merit of an approach that leverages multiple X-ray structures with distinct diverse conformations for MD simulations and machine-learning-based MSM construction to elucidate complex conformational dynamics, and validates the resulting model experimentally with testable biophysical predictions (**Figure 3**). Previously, individual components of our integrative strategy have been employed to study the dynamics of transcriptional activators^48^, kinases^11,12^, and allosteric regulation^13^. However, it is the first time that these diverse approaches are consolidated explicitly with the goal of illuminating conformational dynamics of an enzyme in a comprehensive and feasible manner. Assessment of key computational parameters concluded that we have utilized sufficient diverse seed structures and simulation time for microstate discovery and thus robust construction of conformational landscapes (**Figure 4**). Notably, we relied on a unique computational resource---Folding@home---to collect remarkable six-millisecond simulation data (see Method). Without access to Folding@home, contemporaneous progress on developing *adaptive* Markov state model construction algorithms---where iterative model building guides the collection of additional simulation data^49,50^---will still allow research groups to achieve this feat on local GPU clusters or cloud resources in the near future. Furthermore, the concept of adaptive model construction can be extended to identify which new structural or biophysical data would be valuable in reducing uncertainty^51–53^ and producing refined MSMs. The integrated platform and concept formulated via this work can be readily transformed to explore dynamic conformational landscapes of other proteins.

This work represents the first time that conformational dynamics of a protein methyltransferase has been definitively characterized with atomic details. Strikingly, SETD8 adopts extremely diverse dynamic conformations in apo and SAM-bound states (24 and 10 kinetically metastable macrostates, respectively, **Figure 2**). Interconversions between metastable conformers cover a broad spatio-temporal scale in particular associated with motions of SETD8’s SET-I and post-SET motifs (**Figures 1,5**). In the apo landscape, the general structural features of the X-ray structures of **BC-Inh1**, **BC-Inh2**, **APO**, **BC-SAM**and **TC** (**Figure 1**) are recapitulated by a subset of macrostates (*e.g.* A6 for **BC-Inh1**; A10 for **BC-Inh2**; A12 for **APO**; A20 for **BC-SAM**; A22, A24 for **TC**, 6 of 24 macrostates, **Figure 5**). Such observation indicates that these X-tray structures trapped in the different ligand-binding states are not ligand-induced artifacts but indeed relevant snapshots of hidden conformations of apo-SETD8. Similarly, a few macrostates in the SAM-bound landscape also recapitulate major structural features of the two cofactor-bound X-ray structures (*e.g.* S1, S2, S10 for BC-SAM, S4, S6, S8 for TC, 6 of 10 macrostates, **Figure 5**). Meanwhile, our results also demonstrate that X-ray crystallography alone is insufficient to capture all metastable conformations of SETD8. Remarkably, there is no correlation of overall structural similarity and interconversion rates between metastable conformers. Though the anticipated findings of fast transitions between structurally similar conformers and slow transitions between structurally distinct conformers (*e.g.* microstates within individual satellite macrostates A17–A24 of apo SETD8; S6, S8, and S10 of SAM-bound SETD8, **Figure 5**), we frequently observed fast kinetics of transitions between structurally distinct microstates (*e.g.* microstates within hub-like macrostates A11 and S8; multi-connected states A1–A4, A9, A14, S1 and S3–S5) and *vice versa* (*e.g.* macrostates A22 and A24) (**Figures 2**,**5**). It is thus interesting to examine how other factors such as specific residue contacts and cooperative long range motions of certain structural motifs play roles on interconversion kinetics.

Functional annotation of the landscapes revealed that the SET-I motif adopts diverse conformations (Figure 2), and its overall configuration is a key feature that differentiates the lobes of the dumbbell-like conformational landscape of SETD8. The conformational dynamics within the hub-like macrostate A11 mainly involves motions of the SET-I motif. Two gain-of-function I293G and E292G variants of SETD8 were designed for relaxing distorted configurations of the SET-I motif upon SAM binding (**Figure 3**). These findings argue the functional essentiality of the intrinsically dynamic motions of SET-I motif for SETD8 SAM binding and catalysis. Importance of dynamic conformational modulation of the SET-I motif has also been shown for other SET-domain PKMTs. For instance, the SET domains of MLLs and EZH1/2 alone are catalytically inert but active in the presence of binding partners WDR5-RbBP5-Ash2L-Dpy30 (referred as MLL-WRAD) and EED-Suz12 (referred as PRC2), respectively15. Recent structural evidence implicated that the formation of these complexes regulates the conformational dynamics of the SET-I motif, which is essential for catalysis54,55. Interestingly, this region has also been exploited by cancer-associated mutants of PKMTs. For instance, NSD2’s E1099 is located in its SET-I motif and its E1099K mutant was characterized as a hot-spot cancer mutation with the gain-of-activity of H3K36 methylation56. Additionally, many mutations of PKMTs have been mapped in their SET-I motifs, implicating their potential roles in alternation of function (**Figure S39**, **Table S12**).

In contrast to static X-ray structures, dynamic conformational landscapes greatly facilitated the characterization of cancer-associated SETD8 mutants (**Figure 6**). A significant portion of cancer-associated, loss-of-function SETD8 mutations, though remote from active sites, were revealed to act allosterically through perturbing the SET-I motif and thus catalysis (**Figure 6**). We also discovered significant changes in the connective networks and a dramatic decrease in conformational heterogeneity upon SAM binding (**Figure 2**). This finding highlights how enzyme-ligand interactions reshape conformation landscapes. The conformational landscapes of SETD8 thus provide an unprecedented platform for virtual screening of ligand candidates as inhibitors via exploring different modes of interaction (SAM-competitive, substrate-competitive, covalent or allosteric). Uncovering hidden conformations can thus be essential for developing potent and selective SETD8 inhibitors. The conformations of individual SETD8 microstates can be further explored to derive their thermodynamic, kinetic, and even transition-state parameters in a catalytic cycle. Similar strategies can be generally applied to native or disease-associated PKMTs for functional annotation.

## Acknowledgements

The authors thank for the National Institutes of Health of USA (ML: R01GM096056, R01GM120570; JDC: R01GM121505; JJ: R01GM122749, R01HD088626; YGZ, R01GM126154), National Cancer Institute (ML, JDC: 5P30 CA008748; JJ: R01CA218600), MSKCC Functional Genomics Initiative (ML), the Sloan Kettering Institute (ML, JDC, KAB), Mr. William H. Goodwin and Mrs. Alice Goodwin Commonwealth Foundation for Cancer Research, and the Experimental Therapeutics Center of Memorial Sloan Kettering Cancer Center (ML), and Louis V. Gerstner Young Investigator Award (JDC), K. C. Wong Education Foundation (CL), Chinese Academy of Sciences (CL: XDA12020353), National Natural Science Foundation of China (CL: 81625022 and 81430084), the Tri-Institutional PhD Program in Chemical Biology (RPW and SC), Peer Reviewed Cancer Research Program of the Department of Defense (RPW: W81XWH-17-1-0412) for research supports; the Marie-Josée and Henry R. Kravis Center for Molecular Oncology, and the Molecular Diagnostics Service in the Department of Pathology for the access of tumor mutation data via cBioPortal; Carolina Adura at High Throughput and Spectroscopy Resource Center at The Rockefeller University for the assistance of ITC experiments; Henry Zebroski III and Susan Powell at Proteomics Resource Center at The Rockefeller University for peptide synthesis; the Folding@home project for computational resources; Kanishk Kapilashrami, Josh Fass, Sonya Hanson, Frank Noé, Simon Olsson, and Martin Scherer for insightful discussions or software support. The Structural Genomics Consortium is a registered charity (no. 1097737) that receives funds from AbbVie; Bayer Pharma AG; Boehringer Ingelheim; Canada Foundation for Innovation; Eshelman Institute for Innovation; Genome Canada; Innovative Medicines Initiative (EU/EFPIA) (ULTRA-DD grant no. 115766); Janssen; Merck & Co.; Novartis Pharma AG; Ontario Ministry of Economic Development and Innovation; Pfizer; São Paulo Research Foundation-FAPESP; Takeda; and the Wellcome Trust. The X-ray structure results of **BC-Inh2** and **BC-SAM** are derived from work performed at Argonne National Laboratory, Structural Biology Center (SBC) at the Advanced Photon Source. SBC-CAT is operated by UChicago Argonne, LLC, for the U.S. Department of Energy, Office of Biological and Environmental Research under contract DE-AC02-06CH11357. These experiments were performed using beamline 08ID-1 at the Canadian Light Source, which is supported by the Canada Foundation for Innovation, Natural Sciences and Engineering Research Council of Canada, the University of Saskatchewan, the Government of Saskatchewan, Western Economic Diversification Canada, the National Research Council Canada, and the Canadian Institutes of Health Research. The X-ray experiment of **BC-Inh1** was conducted with NE-CAT beam line 24-ID-E (GM103403) and an Eiger detector (OD021527) at the APS (DE-AC02-06CH11357).

## Author Contributions

M.L., J.D.C, C.L., S.C., and F.M. initialized this project. M.L., J.D.C, and C.L. designed experiments and directed the project. N.B., A.M., J.W., G.B., F.P.-S., J.J., and M.L. developed inhibitors. N.B., A.M., Y.Y., W.T., H.Z., R.S., P.J.B., and J.J. solved X-ray structures. F.M. S.F., H.J., K.C. and C.L. performed initial simulation and analysis. R.P.W. and K.A.B. set up MD simulations. R.P.W. and S.C. conducted computational analysis. S.C., K.Q., H.H., J.W., Y.G.Z., and M.L. designed and conducted biochemical experiments. M.L., J.D.C., S.C., and R.P.W. wrote the manuscript.

## PDB files

6BOZ for **BC-Inh1**, 5W1Y for **BC-Inh2**, 4IJ8 for **BC-SAM**, and 5V2N for **APO**.

## Code and Data Availability

The molecular dynamics datasets generated and analyzed in this study are available via the Open Science Framework at https://osf.io/2h6p4. The code used for the generation and analysis of the molecular dynamics data is available via a Github repository at https://github.com/choderalab/SETD8-materials.

## Online Methods

### Synthesis of inhibitors

Detailed procedures for synthesis and characterization of intermediates are presented in supplemental information. Characterization of the final compounds **MS4138** (**Inh1**) and **SGSS05NS** (**Inh2**): **MS4138** (**Inh1**), ^1^H-NMR (600 MHz, CD_3_OD), δ 7.65 (s, 1H), 7.19 (s, 1H), 6.28 – 6.16 (m, 2H), 5.67 (dd, *J* = 9.0, 3.0 Hz, 1H), 4.43 – 4.33 (m, 2H), 3.99 (br.s, 3H), 3.74 (t, *J* = 6.9 Hz, 4H), 3.62 (br.s, 2H), 3.52 – 3.44 (m, 2H), 3.39 (t, *J* = 6.8 Hz, 2H), 2.15 (br.s, 2H), 2.05 (br.s, 2H), 1.98 (dt, *J* = 13.8, 6.8 Hz, 2H). ^13^C-NMR (151 MHz, CD_3_OD) δ 168.3, 160.4, 155.1, 151.6, 148.6, 136.8, 132.0, 126.8, 105.5, 104.7, 101.2, 66.8, 57.0, 47.4 (two carbons), 40.3, 40.0, 38.0, 29.7, 26.8, 25.6. HRMS calcd for C_21_H_30_N_6_O_3_ + H, 415.2452; found, 415.2444 [M + H]^+^. **SGSS05NS** (**Inh2**), ^1^H-NMR (500 MHz, Chloroform-*d*), δ 8.07 (dd, *J* = 7.1, 1.7 Hz, 1H), 8.02 (dd, *J* = 6.8, 1.6 Hz, 1H), 7.70 – 7.65 (m, 2H), 7.25 – 7.21 (m, 4H), 7.17 – 7.13 (m, 1H), 3.51 (dd, *J* = 6.2, 3.9 Hz, 4H), 2.58 – 2.49 (m, 4H), 2.31 (s, 3H). ^13^C-NMR (151 MHz, chloroform-*d*) δ 182.47, 182.11, 154.17, 136.29, 134.34, 133.29, 132.86, 132.37, 129.36, 128.14, 127.09, 126.91, 126.67, 55.68, 51.37, 46.15. HRMS calcd for C_21_H_20_N_2_O_2_S+ H, 365.1324; found, 365.1331 [M + H]^+^.

### Preparation of SETD8 and its mutants for biochemical assays

Human SETD8 catalytic domain (Uniprot Q9NQR1-1 positions 232-393, SRKSKAELQSEERKRIDELIESGKEEGMKIDLIDGKGRGVITKQFSRGDFVVEYHGDLIEITDAKKREALYAQDPSTGCYMYYFQYLSKTYCVDATRETNRLGRLINHSKCGNCQTKLHDIDGVPHLILIASRDIAAGEELLDYGDRSKASIEAHPWLKH) with an *N*-terminal 6×His tag in pHIS2 vector was overexpressed in *E. coli* Rosetta 2(DE3) in LB medium in the presence of 100 μg/ml of ampicillin. Cells were grown at 37 °C to an OD_600_ of 0.4~0.6 and the expression of SETD8 was induced by 0.4 mM isopropyl-1-thio-*D*-galactopyranoside (IPTG) at 17 °C overnight. Harvested cells were suspended in a lysis buffer (50 mM Tris-HCl, pH=8.0, 25 mM NaCl, 10% Glycerol, 25 mM imidazole) supplemented with EASY pack protease inhibitor (1 tablet/10 mL solution), a tip amount of lysozyme and DNAase I. The mixture was lysed by FrenchPress. SETD8 (aa 232-393) was purified by a Ni-NTA column subjected to a washing buffer (50 mM Tris-HCl, pH=8.0, 25 mM NaCl, 10% glycerol, 25 mM imidazole) and then an eluting buffer (50 mM Tris-HCl, pH=8.0, 25 mM NaCl, 10% glycerol, 400 mM imidazole). The protein was further purified by a Superdex-75 gel filtration column with a buffer containing 25 mM Tris-HCl (pH = 8.0), 200 mM NaCl, and 10% glycerol. The elution fractions were pooled, supplemented with 5 mM of tris(2-carboxyethyl)phosphine (TCEP), and concentrated to about 60 mg/mL for storage at −80 °C. All purification was conducted at 4 °C. The *N*-terminal 6×His SETD8 (aa 232-393) construct was used to measure IC_50_ of SETD8 inhibitors. Plasmids of SETD8 mutants were generated by QuickChange site-directed mutagesis kit (Stragaene) according to manufacturer’s instructions and validated by DNA sequencing. Primer sequences for mutagesis were designed by PrimeX and listed in **Table S2**. SETD8 mutants were expressed and purified as described above for wild-type SETD8.

### Measurement of IC_50_ of SETD8 inhibitors

The IC_50_ of SETD8 inhibitors were measured by a previously reported filter plate assay^31,57^ with some modifications. DMSO stock solutions of SETD8 inhibitors with different concentrations were prepared through series dilution. The final assay mixture (a total volume of 20 μL) contains 300 nM SETD8 protein (*N*-terminal 6×His-taged, amino acid 232-393), 10 μM H4K20 peptide (aa 10-30, prepared by Rockefeller University Proteomics Resource Center, New York, NY), 1.5 μM [^3^H-Me]-SAM (PerkinElmer Life Sciences), and various concentrations of inhibitors in a reaction buffer (50 mM HEPES, pH=8.0, 0.005% Tween-20, 5 μg/mL BSA, 1 mM TCEP and 0.5% DMSO). Prior to each reaction, 10 μL of a reaction mixture containing 2× concentrations of SETD8 and inhibitors was pre-incubated at ambient temperature (22 °C) for 2 hours. 10 μL of another reaction mixture containing 2× concentrations of peptide and [^3^H-Me]-SAM was then added to initialize the reaction. The resulting mixture was allowed to react at ambient temperature (22 °C) for 2 hours. 3×6 μL (total 18 μL) of this mixture were spotted onto 3 wells of MultiScreen_HTS_ PH Filter plate (Millipore) to immobilize ^3^H-labeled peptide. After drying in ambient air overnight, each well was washed 6 times with 200 μL of 50 mM Na_2_CO_3_/NaHCO_3_ buffer (pH=9.2), followed by the addition of 30 μL Ultima Gold scintillation cocktail (PerkinElmer Life Sciences). The plate was sealed and the mixture was further equilibrated for 30 minutes. The immobilized radioactivity of ^3^H-labeled peptide was quantified by 1450 Microbeta liquid scintillation counter. The inhibition curve was generated according to the equation: Percentage of inhibition = [(CPM of no inhibitor control – CPM of a reaction mixture)/(CPM of no inhibitor control − CPM of background)] × 100%. The IC_50_ values were obtained by fitting inhibition percentage *versus* concentrations of inhibitors using GraphPad Prism. Data presented are best fitting values ± s.e‥

### Crystallography

#### BC-Inh1 (6BOZ)

Human SETD8 catalytic domain (amino acid 232-393) with a C343S mutation and an *N*-terminal 6×His tag in pHIS2 vector was overexpressed in *E. coli* BL21-CodonPlus(DE3)-RIL in Terrific Broth medium in the presence of 100 μg/ml of carbenicillin and 30 μg/ml of chloramphenicol. Cells were grown at 37 °C to an OD_600_ of 2.5 and SETD8 expression was induced by 0.3 mM IPTG with a supplement of 1 mM zinc sulfate at 15 °C overnight. Harvested cells were suspended in a lysis buffer (50 mM sodium phosphate, pH=7.5, 0.5 mM NaCl, 5% glycerol) and lysed by microfluidizer. The SETD8 protein (aa 232-393) was purified by a Ni-NTA column. The column was washed by a washing buffer (50 mM sodium phosphate, pH=7.5, 0.5 mM NaCl, 5% glycerol) and the protein was eluted by an eluting buffer (50 mM Tris, pH=8.0, 250 mM NaCl, 250 mM imidazole, 0.5 mM TCEP). *N*-terminal His tag was removed by TEV protease. The protein was further purified by a Superdex 200 (26/600) gel filtration column with a buffer containing 50 mM Tris-HCl (pH=8.0) and 150 mM NaCl. The elution fractions were pooled and supplemented with 0.5 mM of TCEP. All purification steps were performed at 4 °C and in the presence of a protease inhibitor AEBSF (Goldbio).

The purified SETD8 protein sample was mixed with **Inh1 (MS4138)** at a molar ratio of 1:5, and incubated at 4 °C overnight. The solution was then concentrated to about 20 mg/mL and crystallized with the hanging drop vapor diffusion method at 17 °C by mixing equal volume of the protein solution with the reservoir solution (0.1 M HEPES, pH=7.0, 20% (w/v) PEG 6,000, 0.2 M MgCl_2_). SETD8-MS4138 crystals (**BC-Inh1**) were soaked in the corresponding reservoir liquor supplemented with 20% ethylene glycol as cryoprotectant before flash freezing in liquid nitrogen. X-Ray diffraction data were collected at 100K at NE-CAT beamline 24-ID-E of Advanced Photon Source (APS) at Argonne National Laboratory. The data integration and reduction were performed with MOSFLM and SCALA, respectively, from the CCP4 suite^58^. The structures of the SETD8-MS4138 complex were solved by molecular replacement using PHASER software^59^ using the atomic model of the SETD8 catalytic domain (PDB file 4IJ8). The locations of the bound molecules were determined from a Fo-Fc difference electron density map. REFMAC^60^ and phenix.refine^61,62^ were used for structure refinement. Graphic program COOT^63^ was used for model building and visualization. The overall assessment of model quality was performed using MolProbity^64^. Data reduction and refinement statistics are summarized in **Table S1**.

#### BC-Inh2 (5W1Y)

Human SETD8 catalytic domain (amino acid 232-393) with a C343S mutation and an *N*-terminal 6×His tag in pHIS2 vector was overexpressed in *E. coli* BL21 (DE3) V2R-pRARE in Terrific Broth medium in the presence of 50 μg/ml of ampicillin and 50 μg/ml of chloramphenicol. Cells were grown at 37 °C to an OD_600_ of 1.5 and SETD8 expression was induced by 1 mM IPTG at 15 °C overnight. Harvested cells were suspended in lysis buffer (50 mM Tris-HCl, pH=8.0, 300 mM NaCl, 20 mM imidazole, 1 mM phenylmethyl sulfonyl fluoride (PMSF)) and lysed by sonication. SETD8 (aa 232-393) was purified by Ni-NTA column. The column was washed by a washing buffer (50 mM Tris-HCl, pH=8.0, 300 mM NaCl, 20 mM imidazole) and the protein was eluted by an eluting buffer (50 mM Tris-HCl, pH=8.0, 300 mM NaCl, 250 mM imidazole). *N*-terminal His tag was removed by TEV protease. The protein was further purified by a Superdex-75 gel filtration column with a buffer containing 50 mM Tris-HCl (pH=8.0), 100 mM NaCl and 5 mM 1,4-dithiothreitol (DTT). The elution fractions were pooled and concentrated to about 0.7 mg/mL.

The purified SETD8 protein sample was mixed with **Inh2** (**SGSS05NS**) at a molar ratio of 1:3, and incubated on ice for 3 hours until SETD8 was completely covalently modified (confirmed by mass spectrometry). Crystals were initially obtained with a sitting-drop vapor diffusion method at the condition of 0.2 M NaF, 20% w/v polyethylene glycol 3350 by mixing 0.5 uL of this solution with 0.5 uL of the SETD8-**Inh2** solution against 90 uL reservoir buffer at 18 °C. Crystals grow to a mountable size in three days, and soaked in reservoir solution with newly added glycerol (v/v 15%) as a cryoprotectant before mounting. Diffraction data were collected under cooling at beam line 19ID of the Advanced Photon Source and reduced with XDS^65^. Intensities for a 100-degree wedge of the images were merged with POINTLESS/AIMLESS^66^. The structure was solved by molecular replacement with PHASER software^67^ and coordinates from the SETD8-SAM complex (see below). Geometry restraints for the compound were calculated with PRODRG^68^ or, for later stages of refinement, with GRADE^69^, which uses MOGUL^70^. The protein model was automatically rebuilt with ARP/wARP^70,71^. REFMAC^72^ and AUTOBUSTER^73^ were used for restrained refinement^74^. COOT and MOLPROBITY were used for interactive rebuilding and geometry validation, respectively^61,64,75^. Data reduction and refinement statistics are summarized in **Table S1**.

#### BC-SAM (4IJ8)

The conditions for expression and purification of SETD8 (amino acid 232-393 containing a C343S mutation) for crystallography of **BC-SAM** is similar to those of **BC-Inh2** with slight modifications. Purified protein samples were concentrated to about 18 mg/mL, and then mixed with SAM at a molar of 1:10 and incubated on ice for one hour. The sample was crystallized using the sitting drop vapor diffusion method at 18 °C. The crystals of SETD8 in complex with SAM were grown in a condition of 1.08-1.2 M trisodium citrate and 100 mM HEPES (pH = 7.5). SETD8-SAM crystals were soaked in the corresponding reservoir liquor supplemented with 20% ethylene glycol as cryoprotectant before flash freezing in liquid nitrogen. Diffraction images were collected at beam line 08ID of the Canadian Light Source^76^. Diffraction images were processed with the HKL software suite^77^ for early stages of structure determination. For later steps of model refinement, diffraction images were processed with XDS, and intensities further scaled with SCALA^78^. A starting model was obtained from an isomorphous crystal structure, which had been solved by molecular replacement with coordinates from PDB entry 1ZKK^35^. The model was automatically rebuilt with ARP/wARP, manually rebuilt with COOT, and refined with REFMAC. Data reduction and refinement statistics are summarized in **Table S1**.

#### APO (5V2N)

Human SETD8 catalytic domain (amino acid 231-393) with mutations of K297A, K298A, E300A and an *N*-terminal 6×His tag in pET28 vector was overexpressed in Rosetta2(DE3) *E. coli* strain in LB medium in the presence of 50 mg/L kanamycin and 34 mg/L chloramphenicol. The K297A, K298A, E300A mutants were introduced to reduce entropy at the protein surface and thus enhance the ability of apo-SETD8 to crystallize. Cells were grown at 37 °C to an OD_600_ of 0.8 and SETD8 expression was induced by 0.4 mM IPTG at 17 °C overnight. Harvested cells were suspended in lysis buffer containing 25 mM Tris (pH = 7.6), 500 mM NaCl, 0.25 mM TCEP, 0.5 % Triton X-100, and protease inhibitors, and lysed by microfluidizer. SETD8 (aa 231-393) was purified by a Cobalt column. The column was washed by a washing buffer containing 25 mM Tris (pH = 7.6), 500 mM NaCl, 0.25 mM TCEP. The protein was eluted by an eluting buffer containing 25 mM Tris (pH = 7.6), 500 mM NaCl, 200 mM imidazole, and 0.5 mM TCEP. *N*-terminal 6×His tag was removed by TEV protease. The protein was further purified by a Superdex-75 gel filtration column with a buffer containing 20 mM Tris-HCl (pH = 7.0), 100 mM NaCl and 1mM TCEP. The elution fractions were pooled and dialyzed against 20 mM Tris-HCl (pH=7.0), 100 mM NaCl and 1mM TCEP. The peak fractions were pooled, concentrated to 26 mg/ml, immediately frozen as aliquots with liquid nitrogen.

Initial crystal trials were conducted with Takeda California’s automated nanovolume crystallization platform. The purified SETD8 protein sample (26 mg/ml) was crystallized with a sitting drop vapor diffusion method at 20 °C with reservoirs containing 100 mM Tris (pH 8.2-8.8), 30% PEGMME 550, and 5% ethylene glycol. Crystals were soaked in the corresponding reservoir liquor supplemented with 22% ethylene glycol as cryoprotectant before flash freezing in liquid nitrogen. Diffraction data were collected from a single cryogenically protected crystal at the Advanced Photon Source (APS) beamline 23-ID-B at Argonne National Laboratory. Data were reduced using the HKL2000 software package^77^. The structure was determined by molecular replacement with either MOLREP^79^ of the CCP4 program suite utilizing the SETD8 catalytic domain (PDB file 4IJ8) as search model, and refined with the program REFMAC^60^. Several cycles of model building with XtalView^80^ and refinement were performed for improving the quality of the model. Data reduction and refinement statistics are summarized in **Table S1**.

#### Preparation of SAM-free SETD8

SAM-free SETD8 was prepared as described previously^81^. Briefly, the concentrated *N*-terminal 6×His-tagged SETD8 protein (aa 232-393, ~ 60 mg/mL) was diluted with about 1:10 ratio (v/v) by a stripping buffer (25mM Tris-HCl, pH = 8.0, 35 mM KCl, 5% glycerol). Activated charcoal was added into the solution (1:1 w/w ratio of protein versus charcoal). The resulting mixture was incubated for 45 minutes. The charcoal-treated sample was then centrifuged and filtered to afford SAM-free SETD8. All these steps are performed at 4 °C. SAM-free SETD8 mutants were prepared in a similar manner.

#### Isothermal titration calorimetry (ITC)

Dissociation constants of SETD8 with SAM (Sigma-Aldrich) were measured using an Auto-iTC200 calorimeter (MicroCal) at 20 °C. Both SAM and SAM-free SETD8 proteins were dissolved into an assay buffer containing 50 mM HEPES (pH=8.0), 0.005% Tween-20, 5 μg/mL BSA, 0.00125% TFA, and 1 mM TCEP. 2.5 mM SAM was titrated into 125 μM SETD8 through 20 injections. Experimental data were analyzed by Origin 7.0 after correcting the heat generated upon injecting SAM into the assay buffer. Best fits were obtained with a fixed stoichiometry (N=1). Data are shown as mean ± s.e. of at least 3 replicates.

#### Stopped-flow rapid mixing experiment

The binary binding kinetics of SAM to SETD8 (wild-type and mutants) was studied using stopped flow spectrometry (SX20, Applied Photophysics). The slit widths of the entrance and exit of the monochromator were set to 2.0 mm. Equal volume of samples from two 2.5 mL syringes were driven into a 20-μL observation cell to mix at ambient temperature (22 °C), to reach the final concentration of 1 μM SAM-free SETD8 and serial concentrations of SAM (16 μM to 2000 μM) in a mixing buffer containing 50 mM HEPES-HCl (pH = 8.0), 0.005% Tween 20 and 1 mM TCEP. 6–8 shots (drives) were taken for each SAM concentration. Trp fluorescence change was recorded for 1 second upon mixing with an excitation wavelength of 295 nm and a wavelength cutoff emission filter (≥ 320 nm). 10000 data points were collected with Pro-Data SX20 software for each stopped-flow experiment. Data analysis was performed using KinTek Explorer^82^. For the global fitting, the signal traces for all concentrations of SAM were simultaneously fitted to a two-step binding model with an initial binding step followed by the step of further conformational changes: E + SAM = ES= E’S, in which E, ES, and E’S correspond to different states of SETD8. The fluorescence signal was defined as the expression *F* = *a*×[E] + *b*×[ES] + *c*×[E’S] + bkg, in which F is the detected total fluorescence intensity, a, b, and c are fluorescence coefficients of E, ES, and E’S, respectively, and bkg is the background fluorescence intensity. For the calculation of equilibrium constants, the equations of *K*_d1_ = *k_-1_*/*k_1_*, *K*_eq_ = *k_-2_*/*k_2_*, and *K*_d_ = *K*_d1_ × *K*_eq_/(1+ *K*_eq_) were followed. For conventional fittings, the fluorescence data were fitted into eq. 1, in which *F* is the fluorescence intensity, *A_1_* and *A_2_* are the amplitude of the signal changes for fast and slow phases, respectively, *k*_obs_^fast^ and *k*_obs_^slow^ are the observed rate constants for two phases, and *t* is time. The plot of *k*_obs_^fast^ and *k*_obs_^slow^ versus SAM concentrations were fitted with eq. 2 and eq. 3, respectively, where obs obs [S] is the concentration of SAM, *k*_i_ and *k*_-i_ are the association and dissociation rate constants for step i (i = 1 or 2), respectively. For individual rate constants, data are best fitting values ± s.e. from KinTek. Uncertainties of *K*_d1_, *K*_eq_, *K*_d_ are shown as s.e. calculated by the propagation of s.e. from individual rate constants and dissociation constants, respectively.

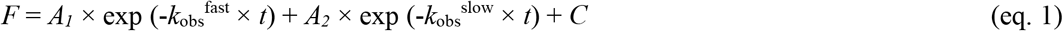

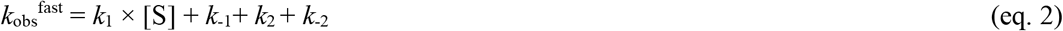

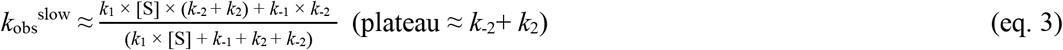

#### Stopped-flow rapid dilution experiment

25 μM SAM-free SETD8 (wild-type and mutants) was pre-mixed with serial concentrations of SAM (1000 μM to 2000 μM) in the mixing buffer and incubated for 10 min at ambient temperature (22 °C). The pre-mixed samples were loaded into a 100 μL syringe, and the mixing buffer was loaded into a 2.5 mL syringe. The two syringes were then driven into the observation cell and mixed to achieve a 1:25 dilution of the pre-mixed samples. The time-dependent fluorescence signal changes were recorded up to 3 s under the same setting as described above for the binding assay. Total of 11333 points were collected with 10000 points for the first 1 s and 1333 points for 1–3 s. Conventional fitting of results was performed using KinTek Explorer following equation: *F*=*A_1_*×exp (-*k*-1×*t*)+*C*, in which *A_1_* is the amplitude of the signal change, *k*_-1_ is the dissociation constant for the first step in rapid quenching experiment, and *t* is time. Signals from different concentrations of SAM are fitted separately, and the average *k*_-1_ is calculated accordingly. Data are best fitting values ± s.e. from KinTek.

#### Methyltransferase assay of cancer-associated SETD8 mutants

The methyltransferase activities of wild-type and cancer-associated SETD8 mutations were characterized by a previously described filter paper assay^31,57^ with some modifications. Briefly, 50 nM SETD8 protein (*N*-terminal 6×His tag, amino acid 232-393, wild-type or mutants), 1.5 μM [^3^H-Me]-SAM, and 30 μM histone H4 peptide (amino acid 10-30) were incubated in a reaction buffer containing 50 mM HEPES (pH=8.0), 0.005% Tween 20, 5 μg/mL BSA, and 1 mM TCEP at ambient temperature (22 °C) for 3 hours. Each reaction mixture was split into 3 aliquots and quenched by spotting on phosphor cellulose (P-81) filter paper, followed by 2-hour air-dry. The dried filter paper was then washed 5 times with 50 mM Na_2_CO_3_/NaHCO_3_ solution (pH=9.2). The washed filter paper was then transferred into a scintillation vial, well mixed with 0.5 ml ddH_2_O and 5 ml UltimaGold^TM^, and analyzed by a Liquid Scintillation Analyzer (Perkin Elmer Tri-Carb 2910 TR). The methyltransferase activities of SETD8 mutants relative to that of wild-type SETD8 were calculated with the following equation: Percentage of relative activity = [(CPM of mutant − CPM of background)/(CPM of wild type − CPM of background)]×100%. Data are presented as mean ± s.d. of 3 replicates.

### Apo-SETD8 simulations

#### Preparation of molecular dynamics (MD) simulations

All-atom models of the 162-residue SET-domain-containing apo-SETD8 fragment (amino acid 232-393, corresponding to the catalytic domain used in our biochemical experiments) were prepared using Ensembler 1.0.5^83^ with default parameters unless otherwise specified. Ensembler automatically corrects sequence variations and models in missing atoms^83^. To prepare apo protein models with diverse conformations for simulation, the crystal structures of **BC-Inh1** (6BOZ), **BC-Inh2** (5W1Y), **BC-SAM** (4IJ8), **APO** (5V2N), and **TC** (1ZKK, 2BQZ, 3F9W, 3F9X, 3F9Y, 3F9Z) together with four structural chimeras were used as templates for UCSF MODELLER 9.16^84^ (see **Table S3** for details). The structural chimeras were constructed with MDTraj 1.7.2^85^ by combining the C-flanking domain (residues 377-393) with the rest of the protein from different crystal structures (see SI for details). Using OpenMM 6.3.1^86^, protonation states appropriate for pH 7 were assigned with openmm.app.modeller, which uses intrinsic p*K_a_* values to determine the most likely ionization states of individual residues but ensures all models are created with the same protonation and tautomeric state so they can be analyzed collectively. The protein was then energy-minimized and relaxed with 100 ps of implicit solvent dynamics using the OpenMM Langevin integrator with a 2 fs timestep and a 20 ps^−1^ collision rate in the NVT ensemble (T = 300 K). All covalent bonds involving hydrogen were constrained. The protein was then solvated with water in a rectilinear box with 1 nm padding on all sides of the protein, and neutralized with a minimal amount of NaCl. All available chains in the template crystal structures were modeled separately (see SI for details), resulting in 30 simulation-ready structures (representing 9 distinct conformers) solvated by an equal number of water molecules (35,200 atoms total). These structures were equilibrated for 5 ns in the NpT (p = 1 atm, T = 300 K) ensemble. Pressure was controlled by a Monte Carlo molecular-scaling barostat with an update interval of 50 steps. Non-bonded interactions were treated with the Particle Mesh Ewald method^87^ using a real-space cutoff of 0.9 nm and relative error tolerance of 0.0005, with grid spacing selected automatically. These simulations were subsequently packaged as seeds for production simulation on Folding@home^88^. For all simulations, the parameter files included in the OpenMM 6.3.1 distribution^86^ were used for the Amber ff99SB-ILDN force field^89^, the GBSA-OBC2 implicit solvent model^90^ (for implicit refinement), the TIP3P rigid water model^91^ (for explicit equilibration and production), and the adapted Aqvist (Na^+^)^92^ and Smith & Dang (Cl^−^)^93^ parameters for NaCl. Default parameters were used unless noted otherwise.

#### Folding@home simulations

The simulation seeds representing 9 distinct conformers (30 distinct structures derived from multiple chains in each PDB structure, see SI for details) were used to initiate massively parallel distributed MD simulations on Folding@home^88^. Production simulations used the same Langevin integrator as the NpT equilibration described above, except that the Langevin collision rate was set to 1 ps^−1^ to provide realistic heat exchange with a thermal bath while minimally perturbing dynamics. In total, 5,020 independent MD simulations were generated on Folding@home^88^: 600 simulations were produced from each seed conformation prepared from the five crystal structures, and 500 or 510 simulations for each seed conformation prepared from the four structural chimeras. At least 500 MD trajectories were produced for each seed conformation. 99.1% of the generated trajectories (4,976 trajectories) successfully reached 1 μs each (see **Figure S16** for length distribution histogram), resulting in 5.058 ms of aggregate simulation time and 10,115,617 frames. This amount of simulation time corresponds to ~ 231 GPU-years on an NVIDIA GeForce GTX 980 processor. Conformational snapshots (frames) were stored at an interval of 0.5 ns/frame for subsequent analysis. Prior to data analysis, the first 50 frames (25 ns) of each trajectory were discarded to allow the trajectories to relax away from their initial seed conformations. On initial analysis of the RMSDs of the trajectories to their starting frames, one trajectory showed the protein unfolding and was removed from the dataset. The resulting final dataset contained 5,019 trajectories, 4.931 ms of aggregate simulation time, and 9,862,657 frames. This trajectory dataset without solvent is available via the Open Science Framework at https://osf.io/2h6p4. The code used for the generation and analysis of the molecular dynamics data is available via a Github repository at https://github.com/choderalab/SETD8-materials.

#### Optimal hyperparameter selection for featurization and tICA

To select the optimal featurization of the data for subsequent Markov state model (MSM) analysis, we used variational scoring^94–97^ combined with cross-validation^38^ to evaluate model quality, consistent with modern MSM construction practice^38^. To evaluate a large set of hyperparameters to achieve optimal featurization, a reduced dataset subsampled to 5 ns/frame intervals (986,464 frames, 10% of the dataset) was used for computational feasibility. The following trajectory featurization choices were assessed: *a*) all residue–residue distances (calculated as the closest distance between the heavy atoms of two residues separated in sequence by at least two neighboring residues) that cross a 0.4 nm contact threshold in either direction at least once (yielding 6,567 of 12,720 total residue-residue distances); *b*) a transformed version of (a) used by MSMBuilder^98^ to emphasize short-range distances in the proximity of residue-residue contact via eq. 4, with *steepness* = 5 nm^−1^ and *center* = 0.5 nm; *c*) backbone (phi, psi) and sidechain (chi1) dihedral angles, with each angle featurized as its sine and cosine (yielding 920 total features).

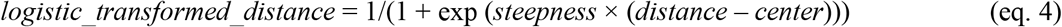

To identify the optimal featurization, we used a 50:50 shuffle-split cross-validation scheme to evaluate various model hyperparameters while avoiding overfitting. In this scheme, subsets of the groups of trajectories initiated from the same conformation (RUNs – see **Table S3** for further explanation) are randomly split into training and test sets of 2,509 and 2,510 trajectories respectively, using scikit-learn 0.9.199. All further steps until scoring were conducted by fitting the model to the training set only, then transforming the test set according to this model. Scoring was based on the sum of the top 10 squared-eigenvalues of the transition matrix (rank-10 VAMP-2^96^). Model scores are reported below as means with standard deviations over five shuffle-splits.

To evaluate each featurization choice, the data were projected into a kinetically relevant space using tICA^100^, retaining all tICs, at lag times of either 5 or 50 ns, with either kinetic^101^ or commute mapping^102^. Each of the tICA outputs was discretized using k-means clustering into 50, 100, 500, or 1000 microstate clusters (see **Table S4** for the summary of options assessed). Featurization was performed using MDTraj 1.8 and PyEMMA 2.4^103^, tICA was performed with PyEMMA 2.4^103^, and clustering was performed with PyEMMA 2.5.1^103^. MSMs at a lag time of 50 ns were constructed with PyEMMA 2.5.1^103^ using discrete microstate trajectories from the training set and scored on the test set trajectories. To obtain standard deviations indicative of out-of-sample model performance, this shuffle-split model evaluation procedure was repeated 5 times with different random divisions of the dataset into training and test sets (see SI for results). The data showed (**Figure S17, S18**) that the four individual models with highest average scores were featurized with dihedral angles (featurization *c;* scores: 9.68 (SD = 0.05), 9.68 (SD = 0.05), 9.63 (SD = 0.03), 9.62 (SD = 0.02)), while the highest median score over all models was the residue-residue distance featurization (the median score of 8.20 (mean = 7.98, SD = 1.11) for featurization *a*; 8.06 (mean = 7.49, SD = 2.07) for featurization *c*; 6.99 (mean = 6.47, SD = 2.13) for featurization *b*). Therefore, both featurizations *a* and *c* were further evaluated on the full dataset to determine the optimal model. For both featurizations, commute mapping resulted in significantly higher scores (**Figure S19, S20**) than kinetic mapping, hence commute mapping was used for the full dataset. The shorter tICA lag time (5 ns) was used for the full dataset, as there was no significant difference in scores between 5 and 50 ns (**Figure S19, S20**).

#### Final featurization and microstate number selection

To determine the optimal number of microstates, we again used variational scoring^94–97^ combined with cross-validation^38^ to evaluate model quality. The full dataset (4.931 ms, 0.5 ns/frame, 9,862,657 frames) was separately featurized with the top-scoring feature sets: 6,567 distances (featurization *a* above) and 920 dihedral angles (featurization *c* above). As for the featurization selection, we used the 50:50 shuffle-split cross-validation scheme, using the same 5 data splits. All further steps until scoring were conducted by fitting the model to the training set only, subsampled to 5 ns/frame intervals for computational feasibility, then transforming the full training set and the test set according to this model. Data were projected into the tICA space using a lag time of 5 ns. The tICs were scaled by commute mapping, with subsequent clustering operations using a sufficient number of tICs necessary to explain 95% of the total kinetic content. The tICA outputs were discretized using k-means clustering into 100, 500, 1000, 2000, 3000, 4000, or 5000 microstate clusters (see **Table S5** for the summary of options assessed). Featurization was performed with MDTraj 1.8 and PyEMMA 2.4^103^, tICA was performed with PyEMMA 2.4^103^, and clustering was performed with PyEMMA 2.5.1^103^. MSMs were constructed with PyEMMA 2.5.1^103^ from the discrete trajectories of the training set using a lag time of 50 ns and subsequently scored on the test set, using the rank-10 VAMP-2^96^ score. The highest scoring model (**Figure S21**) had dihedral features (featurization *c* above) and 100 microstates (VAMP-2 = 9.25 (SD = 0.32)). tICA and k-means clustering were refitted to the full dataset subsampled to 5 ns/frame intervals for computational feasibility. Keeping the number of tICs necessary to explain 95% of the total kinetic content resulted in 466 tICs used for k-means clustering. The full dataset was then transformed to give the final discretized trajectories at 0.5 ns/frame intervals. Checking the convergence of the implied timescales validated the choice of the MSM lag time (**Figure S22**). The Chapman-Kolmogorov test^104^ was then conducted on the MSM to validate the self-consistency of the model (**Figure S23**). To aid structural interpretation, the 10 frames closest to each of the 100 microstate cluster centers were extracted from the dataset.

#### Coarse-graining to kinetically metastable macrostates

To coarse-grain the MSM into a small number of kinetically metastable macrostates, a Hidden Markov Model (HMM) was constructed from the discrete trajectories of the optimal model above using PyEMMA 2.5.1^103^. Increasing numbers of macrostates were explored and interpreted structurally by assigning the 10 frames closest to each of the 100 microstate cluster centers to the macrostate to which they had the largest fractional membership. We chose the minimal number of macrostates that achieved increasing structural separation of the distinct SET-I and post-SET motif configurations and hence constructed a 24-macrostate HMM. The resulting HMM provides both a macrostate-to-macrostate transition matrix and a fractional membership of each microstate to each kinetically metastable macrostate. Checking the convergence of the HMM implied timescales further validated the choice of the MSM/HMM lag time (**Figure S24**). To preserve kinetic relationships between macrostates in a two-dimensional representation, the log-inverse fluxes between all pairs of macrostates (calculated using the third power of the transition matrix to eliminate sparsity) were embedded in two dimensions using iterative multidimensional scaling (MDS)^105–107^ with scikit-learn 0.9.1^99^. MDS was repeated at least 50 times with random initializations and the projection that leads to a figure with the fewest crossings of inter-state flux arrows was selected. To aid structural interpretation, the 10 frames closest to each of the 100 microstate cluster centers were assigned to the macrostate to which they had the largest fractional membership. The RMSDs of the macrostates to the homology models derived from all 5 crystal structures generated by Ensembler were calculated by averaging the RMSDs of all 10 frames in each microstate, then for each macrostate taking the mean over all microstates weighted by the HMM observation probabilities. The RMSDs of the 10 frames in each microstate were calculated with C-alpha atoms only, after first superposing each frame onto the reference structure using only the C-alpha atoms of the conformationally homogenous SET motifs (residues 257-290 and 327-376). To quantify the structural diversity of each macrostate, a sample of 100 frames was drawn from each macrostate with probabilities for each frame given by the observation probability of the frame’s microstate from the given macrostate divided by the total number of frames in the frame’s microstate. The C-alpha RMSD (after superpose of the SET motifs only) of each frame versus all other 99 frames was then calculated, and the minimum average RMSD over all 100 reference frames was reported (**Table S7**)

### SAM-bound SETD8 simulations

#### Preparation of molecular dynamics (MD) simulations

All-atom models of the same 162-residue SETD8 fragment in complex with SAM were prepared in a similar manner as apo-SETD8 except that a manual pipeline was used instead of Ensembler. Briefly, two available cofactor-bound crystal structures were used to generate two seed structures for simulation: 4IJ8 (the crystal structure of the binary complex of SETD8 with SAM) and 2BQZ (the crystal structure of the tertiary complex of SETD8 with SAH and a methylated H4K20 peptide). Among the available tertiary complex structures (1ZKK, 2BQZ, 3F9W, 3F9X, 3F9Y, 3F9Z), 2BQZ was selected for MD simulations because of the following conditions met simultaneously: no mutations present, minimum number of missing residues requiring modeling (1), methylated lysine resolved on the histone peptide (for future simulations of the tertiary complex). Protein chains “A” of both structures were used. Mutations in 4IJ8 were corrected to the reference sequence, and missing protein residues and atoms were added using PDBFixer 1.3^86,108^. To replace SAH with SAM in the 2BQZ model, the coordinates of SAM were copied from 4IJ8, where all SAM atoms were resolved, after aligning the common atoms in SAM and SAH with MDTraj 1.7.2^85^. The peptide and SAH were then removed from 2BQZ. Using OpenMM 7.0.1^109^, protonation states appropriate for pH 7 were assigned with openmm.app.modeller. SAM was modeled in the +1 cationic form at its sulfonium center and the zwitterionic form at its α-amino acid moiety. GAFF force field parameters^110^ and AM1-BCC^111^ charges were assigned using Antechamber^112^, with missing parameters assigned using Antechamber’s ParmChk2. The SAM parameter files were then converted from the Amber format to the OpenMM XML format using the conversion script distributed with the openmm-forcefields package^113^. The systems were solvated in rectilinear water boxes with 1 nm padding and neutralized with a minimal amount of NaCl. This resulted in the final systems containing 34556 atoms (system prepared from 4IJ8) and 35588 atoms (system prepared from 2BQZ). These were energy-minimized and relaxed for 1 fs in the NVT (T = 10 K) ensemble using the OpenMM Langevin integrator with a 0.01 fs timestep, 91 ps^−1^ collision rate. Nonbonded interactions were treated with the reaction field method only during minimization (due to its increased stability over PME when steric clashes needed to be resolved following introduction of mutations)^114^ at a cutoff of 0.9 nm. The systems were then equilibrated for 10 ns in the NpT (p = 1 atm, T = 300 K) ensemble using the OpenMM Langevin integrator, the PME nonbonded method, a Monte Carlo molecular-scaling barostat with anh updated interval of 25 steps, and packaged with OpenMM 6.3.1^86^ as seeds for production simulation on Folding@home^88^. All other force field parameters and simulation settings were as previously described for apo-SETD8.

#### Folding@home simulations

In total, 1,000 independent MD simulations were generated on Folding@home: 500 each for the two seed structures prepared above. Simulations employed the same settings as for NpT production of apo-SETD8. 99.7% of the generated trajectories (997 trajectories) successfully reached 1 μs each (see **Figure S25** for length distribution), resulting in 1.003 ms of aggregate simulation time and 2,005,945 frames. This amount of simulation time corresponds to ~ 46 GPU years on an NVIDIA GeForce GTX 980 processor. Prior to data analysis, the first 25 ns of each trajectory were discarded to allow the trajectories to relax away from the initial equilibrated configurations. One trajectory was shorter than the length being discarded and was removed from the dataset. The resulting final dataset contained 999 trajectories, 0.978 ms of aggregate simulation time, and 1,955,965 frames. This trajectory dataset without solvent is available via the Open Science Framework at https://osf.io/2h6p4. The code used for the generation and analysis of the molecular dynamics data is available via a Github repository at https://github.com/choderalab/SETD8-materials.

#### Coarse-graining to kinetically metastable macrostates

To construct a Hidden Markov model of SAM-bound SETD8, the full dataset (0.978 ms, 0.5 ns/frame, 1,955,965 frames) was featurized using the final model generated from apo-SETD8 (featurization *c*, backbone and sidechain dihedral features). The data were projected into the tICA space derived from the apo-SETD8 simulations and assigned to the 100 k-means microstates of apo-SETD8. The SAM-bound SETD8 trajectories populated 67 of the 100 microstates of apo-SETD8. An MSM with a 50 ns lag time was constructed, and the Chapman-Kolmogorov test^104^ was conducted to validate the self-consistency of the model (**Figure S27**). Finally, a Hidden Markov model (HMM) was constructed for a 50-ns lag time using 10 macrostates (the minimal number of macrostates to achieve increasing structural separation between distinct SET-I and post-SET configurations was chosen in the same way as for apo-SETD8). As for apo-SETD8, log-inverse fluxes between macrostates were used to construct a two-dimensional representation, and the 10 frames closest to the microstate cluster centers were assigned to the macrostate to which they had the highest fractional membership to aid structural interpretation. Prior to visualization, frames were re-imaged with MDTraj 1.8^85^ to ensure SETD8 was centered and the SAM ligand was in the same unit cell. Further, as for apo- SETD8, macrostate C-alpha RMSDs to the homology models derived from all 5 crystal structures were calculated by weighted mean over microstate average RMSDs, and structural diversity was quantified by the reference frame with the minimum average RMSD of each macrostate.

### Cancer-associated mutant apo-SETD8 simulations

#### Preparation of molecular dynamics (MD) simulations

All-atom models of the same 162-residue SETD8 fragment with each of 24 cancer-associated single mutations (including 1 deletion giving a 161-residue fragment) identified from the cBioPortal for Cancer Genomics^42–44^ were prepared in an analogical way to apo-SETD8 using Ensembler 1.0.5. The mutants prepared are summarized in **Table S11**(#1–21, 23–25). As we aimed to gain a direct interpretation of the influence of these mutations on the enzymatic activity of SETD8, only a single chain of the TC structure was used as the template. To choose the particular chain out of the 18 TC chains used in the apo-SETD8 simulations (**Table S3**), the homology models of all the TC chains generated by Ensembler were projected into the apo-SETD8 tICA space using PyEMMA 2.4^103^. The distances between the points in the tICA space were then calculated with SciPy 1.0 and chain “A” of the 1ZKK structure, which had the smallest average distance to all others, was selected. The modeling procedure was the same as for apo-SETD8, except the appropriately mutated sequences were passed to Ensembler^83^. Briefly, homology models were created by UCSF MODELLER 9.16^84^, protonation states appropriate for pH 7 were assigned with OpenMM 6.3.1^86^, the models were then energy-minimized and relaxed with 100 ps of implicit solvent dynamics. The proteins were then solvated in rectilinear boxes with 1 nm padding and neutralized with a minimal amount of NaCl. This resulted in the final systems containing between 35,185 and 35,208 atoms. These were equilibrated for 5 ns and packaged as seeds for production simulation on Folding@home. All force field parameters and simulation settings were as previously described for wild type apo-SETD8.

#### Folding@home simulations

In total, 960 simulations were generated on Folding@home: 40 for each of the mutants. Simulations employed the same settings as for NpT production of wild type apo-SETD8. 99.7% of the generated trajectories (957 trajectories) successfully reached 1 μs each (see **Figure S29**for length distribution), resulting in the final aggregate dataset of 0.966 ms and 1,931,849 frames. This amount of simulation time corresponds to ~ 44 GPU years on an NVIDIA GeForce GTX 980 processor. This trajectory dataset without solvent is available via the Open Science Framework at https://osf.io/2h6p4. The code used for the generation and analysis of the molecular dynamics data is available via a Github repository at https://github.com/choderalab/SETD8-materials.

#### Contact map analysis

Prior to data analysis, the first 750 ns of each trajectory were discarded to allow for successful metastable transitions out of the wild type kinetic basin. For the remaining frames of each mutant, all residue-residue distances (calculated as the closest distance between the heavy atoms of two residues) for which the two residues are separated in sequence by at least two other residues (yielding 12,720 residue-residue distances) were calculated with MDTraj 1.8^85^. These were converted into binary contact maps by replacing all distances below the 0.4 nm contact threshold with 1’s and all other distances with 0’s, and casting into a square-form matrix. These were then averaged over all frames of each mutant, yielding one contact map for each mutant. In the same way, the wild type contact map was calculated using the 60 wild type apo-SETD8 trajectories started from chain “A” of the 1ZKK structure (**Table S3**). The wild type contact map was then subtracted from the mutant contact maps to generate one absolute contact change map for each mutant. In the one case of amino acid deletion, all contact changes corresponding to that residue position were set to zero. Relative contact changes were also calculated by dividing the absolute contact change value by the wild type contact value and taking the modulus of the result. The result was set to zero where the wild type value was zero. Contacts with absolute changes of more than 0.2 and relative changes of more than 3 were selected for further structural annotations.

#### Extraction of hypothetical new conformations

For each mutant, of the contacts that showed an absolute fractional change of more than 0.2 or more negative than −0.2, up to 5 contacts with the most positive changes (“positive contacts”) and up to 5 contacts with the most negative changes (“negative contacts”) were noted. For each trajectory (after discarding the first 750 ns) of a given mutant, every 10th (every 20th if more than 7,500 such frames were present) of the frames that had all “positive contacts” present or none of the “negative contacts” present was extracted. These frames were manually inspected using PyMOL 1.8.4^115^ against the representative conformations of the wild type apo-SETD8 microstates (the 10 frames closest to each of the microstate cluster centers) and frames that displayed similar SET-I and post-SET motif configurations to any of the wild type conformations were discarded. For the remaining frames, the C-alpha RMSDs to all frames of the wild type apo-SETD8 dataset subsampled to 5 ns/frame were calculated using MDTraj 1.8^85^ and the wild type frames with the lowest RMSD to each mutant frame were extracted. The mutant frames were manually inspected using PyMOL 1.8.4^115^ against the extracted wild type frames and further mutant frames similar to the wild type frames were discarded. The remaining frames for each mutant were clustered based on the manual inspection and their C-alpha RMSDs to all frames of the given mutant (without discarding the first 750 ns of each trajectory) were calculated using MDTraj 1.8^85^. For each cluster of the hypothetical new conformations, every 10th of all mutant frames with RMSDs below the 0.3 nm, 0.35 nm, and 0.4 nm thresholds to any of the cluster frames were extracted and manually inspected in PyMOL 1.8.4^115^. The 0.3 nm threshold gave good structural similarities and only a small number of false positives (frames that were not sufficiently similar to the originally chosen hypothetical new conformations) were discarded, while the other two thresholds introduced too many false positives. Hence the remaining frames extracted at the 0.3 nm threshold were taken as the final clusters of hypothetical new conformations.

#### Calculation of microstate coverage

To quantify how the diversity of starting conformations influences the number of microstates observed out of the total of 100, the apo-SETD8 discrete trajectories were split into nine sets corresponding to their starting conformations. All logical relations between the sets were generated and the numbers of microstates explored in each intersection were counted in order to produce Venn diagrams of microstate coverage. Analogically, the SAM-bound SETD8 discrete trajectories were split into two sets and the microstate coverage was evaluated. Further, to quantify how the number and the length of trajectories influence the number of microstates observed in addition to the diversity of starting conformations, all combinations of all possible lengths of the five apo-SETD8 sets started from crystal structures were enumerated. Appropriate sets out of the four originating from structural chimeras were added to those combinations which contained the appropriate SET-I and post-SET motif configurations for the formation of those chimeras. Also, if a combination contained either of the BC-Inh1 or BC-Inh2 sets (SET-I configuration I1), and the BC-SAM set (post-SET configuration P1), the TC set (configurations I1-P1) was added as it could be generated as a structural chimera. For all combinations that resulted, the microstate coverage was assessed at all trajectory numbers between 0 to all trajectories in the combination at intervals of 50 trajectories, and simultaneously at all maximum trajectory lengths between 0 to the length of the longest trajectory in the combination at intervals of 50 ns. The desired number of trajectories was randomly drawn from all trajectories in the combination without replacement and the trajectories were trimmed to the desired maximum length. The number of microstates observed was then calculated. This was repeated five times with different draws of trajectories and the results of the five draws were averaged. Analogically, the microstate coverage at increasing trajectory numbers and trajectory lengths was evaluated for the two SAM-bound SETD8 sets.

